# Human calpain-3 and its structural plasticity: dissociation of a homohexamer into dimers on binding titin

**DOI:** 10.1101/2024.02.28.582628

**Authors:** Qilu Ye, Amy Henrickson, Borries Demeler, Vitor Hugo Balasco Serrão, Peter L. Davies

## Abstract

Calpain-3 is an intracellular Ca^2+^-dependent cysteine protease abundant in skeletal muscle. Its physiological role in the sarcomere is thought to include removing damaged muscle proteins after exercise. Loss-of-function mutations in its single-copy gene cause a dystrophy of the limb-girdle muscles. These mutations, of which there are over 500 in humans, are spread all along this 94-kDa multi-domain protein that includes three 40+-residue sequences (NS, IS1, and IS2). The latter sequences are unique to this calpain isoform and are hypersensitive to proteolysis. To investigate the whole enzyme structure and how mutations might affect its activity, we produce the proteolytically more stable 85-kDa calpain-3 ΔNS ΔIS1 form with a C129A inactivating mutation as a recombinant protein in *E. coli*. During size-exclusion chromatography, this calpain-3 was consistently eluted as a much larger 0.5-MDa complex rather than the expected 170-kDa dimer. Its size, which was confirmed by SEC-MALS, Blue Native PAGE, and AUC, made the complex amenable to single-particle cryo-EM analysis. From two data sets, we obtained a 3.85-Å reconstruction map that shows the complex is a trimer of calpain-3 dimers with six penta-EF-hand domains at its core. Calpain-3 has been reported to bind the N2A region of the giant muscle protein titin. When this 37-kDa region of titin was co-expressed with calpain-3 the multimer was reduced to a 320-kDa particle, which appears to be the calpain dimer bound to several copies of the titin fragment. We suggest that newly synthesized calpain-3 is kept as an inactive hexamer until it binds the N2A region of titin in the sarcomere, whereupon it dissociates into functional dimers.

## Introduction

The calpain superfamily is comprised of Ca^2+^-dependent, intracellular, cysteine proteases (1–3). These multidomain enzymes are tightly regulated and perform limited cleavage of their target proteins to help initiate cellular processes including cell motility, signal transduction, cell cycle progression, apoptosis, and gene expression. Abnormal regulation of calpain proteases has been correlated with various diseases such as neurodegenerative disorders, various cancers, cardiovascular disorders, muscular dystrophy, metabolic diseases, and inflammation among others (4–6). In mammals the calpain family is composed of 15 gene products, nine of which are ubiquitously expressed while six have been found to be tissue-specific based on their gene expression profiles and physiological roles (7). Calpain-3 is in the latter category and is predominantly expressed in skeletal muscle (8). Soon after the discovery of this muscle-specific family member, inactivating mutations in the calpain-3 gene (*CAPN3*) were shown to cause the autosomal recessive form of limb girdle muscular dystrophy type R1 (LGMDR1) (9). The physiological functions of calpain-3 are thought to include the natural turnover of muscle proteins (10, 11), regulation of calcium release and recruitment of binding partners in skeletal muscle (12), and remodelling of the sarcomere upon exercise stress, especially after eccentric exercise (13, 14). However, the precise mechanism by which calpain-3 functions in the physiology of healthy muscle and LGMDR1 pathogenesis remains unclear.

Calpain-1 and -2 are designated as “conventional calpains” and were the first members of the calpain superfamily to have been purified and characterized (1). Conventional calpains have a heterodimeric architecture, composed of a catalytic large 80 kDa subunit and a regulatory small 28 kDa subunit, both of which are required for function (15). The catalytic large subunit contains an N-terminal anchor helix and four domains: the calpain-type cysteine protease core domains PC1 (containing the catalytic cysteine) and PC2 (containing histidine and asparagine, which together form a catalytic triad upon calcium binding) (16), the calpain-type beta-sandwich (CBSW) domain, and the penta-EF-hand (PEF(L), where L indicates large subunit) domain (3). The small subunit of calpain contains a glycine-rich region and a PEF(S) (where S indicates small subunit) domain that serves a regulatory function. Each PEF(L) and PEF(S) domain contain five EF-hand α-helical motifs which fold together to form pairs: EF1 with EF2, EF3 with EF4, while EF5 is unpaired. The EF5 from each of PEF(L) and PEF(S) domains pair up with each other to form a heterodimer. This interaction is stabilized through a short antiparallel beta-sheet, with contributions from some hydrophobic interactions (17–20). The same 5^th^ EF-hand pairing of the homo- and heterodimeric PEF domains takes place whether Ca^2+^ is present or absent.

The domain architecture of calpain-3 is similar to that of the large subunit of conventional calpains being comprised of the protease core domains PC1 and PC2, the CBSW domain, as well as a C-terminal PEF domain (21). A sequence comparison indicates that calpain-3 shares approximately 51% and 54% sequence identity with the large subunit in conventional calpain-1 and calpain-2, respectively, but contains three additional sequences that are unique to calpain-3. These are an N-terminal sequence (NS) at the very N-terminal end, preceding PC1; an insertion sequence 1 (IS1) located in protease core domain PC2; and an insertion sequence 2 (IS2) situated between the CBSW domain and the PEF domain (22, 23). These three additional regions confer unique physiological functions to calpain-3 distinct from the conventional calpains but also predispose it to rapid proteolysis. The half-life of native calpain-3 *in vitro* is less than 10 min, due to autolysis of the NS and IS1 regions (24). This makes purification of calpain-3 from skeletal muscle extremely difficult and is a barrier to studying the natural enzyme.

The NS sequence comprises ∼50 residues that are rich in proline and glycine residues, thus implying an unstructured region (25). The 48-residue IS1 sequence is also largely unstructured but with a short central region that is predicted to be alpha helix (26). The sequences and flanking sequences of NS and IS1 contain several proteolytic cleavage sites that are highly sensitive to exogenous proteases such as trypsin and chymotrypsin. IS1, however, is particularly prone to autoproteolysis (27) because it occupies the catalytic cleft of the protease core (28, 29) and the main autoproteolytic site is immediately adjacent to the active site cysteine. This suggests that IS1 functions as an internal propeptide to help regulate calpain-3 proteolytic activation (26, 28).

The second insertion sequence, IS2, lies between the upstream CBSW domain and the downstream PEF domain. Deletion of the IS2 sequence increases both levels and stability of calpain-3 in the cytosol, suggesting that the IS2 sequence may also contain internal cleavage site(s) associated with the rapid degradation of calpain-3 (23). Further characterization of calpain-3 has demonstrated that the IS2 region can interact directly with the N2A and M-line regions of titin, a gigantic protein in myofibrils, implying that titin may be involved in regulating calpain-3 stability and activity (30–32) through IS2.

Unlike the conventional calpains -1 and -2, calpain-3 is incapable of forming a heterodimer with the ubiquitously expressed calpain small subunit (21, 33). Biochemical and biophysical studies confirmed that recombinantly expressed PEF domain of calpain-3 forms a homodimer in strong preference to a heterodimer with the small subunit PEF domain (34). This was confirmed by the X-ray crystallography determination of the homodimerized PEF domain of calpain-3 (35). As expected, the 5^th^ EF-hand motifs of the PEF domains are paired up, and there is a Ca^2+^ in both of these motifs. Using size-exclusion chromatography, recombinant full-length inactive calpain-3 (C129S) eluted as a putative homodimer (33). More recently an inactive calpain-3 expressed in COS7 cells has been reported by Hata *et al.* to form a homotrimer (36). In our attempts to produce recombinant calpain-3 for structural studies we observed that the enzyme eluted from SEC at an apparent molecular weight larger than expected for a globular dimer. Our model for the dimer (35) indicated that the protein would be elongated and might have a larger apparent molecular weight because of its asymmetry.

To clarify calpain-3’s oligomerization state and work towards an understanding of how LGMDR1 mutations affect the workings of the enzyme, we have used an *E. coli* expression system to produce a stable recombinant inactive calpain-3 variant (C129A) with truncated NS and IS1 regions. Unexpectedly, this inactive full-length calpain-3 (C129A) ΔNS&IS1 largely exists as a homohexamer. Further analysis using single particle cryo-EM indicated that this protease forms a trimer of dimers and that the dimerization interaction is distinctly different from that seen in the crystal structure of the calpain-3 PEF domain homodimer. We also report that the homodimer is split into dimers upon binding to titin’s N2A region and suggest that the oligomer is a way of sequestering calpain-3 until it can bind productively to the sarcomere.

## Experimental procedures

### Cloning and expression of recombinant calpain-3

To avoid rapid proteolysis of recombinant calpain-3 produced in *E. coli*, a construct was made from which 45 residues of NS (P^2^ – I^46^) and 48 residues of IS1 (D^268^GTNM-----RPTR^315^) were deleted. In addition, the catalytic cysteine of the active site (Cys129) was changed to Ala129 to eliminate autoproteolytic activity. Human calpain-3 isoform_1 (NM_000070.3) was used as a template from which GeneArt (Thermo Fisher Scientific) synthesized the modified gene following codon optimization. The synthesized DNA was inserted into a pET24a vector such that the expressed product would have a C-terminal hexa-histidine tag under kanamycin selection. Positive clones were verified by DNA sequencing and transformed into the BL21PLySs (DE3) strain of *E. coli*. For the co-expression of calpain-3 with titin-I81-I83, the human titin N2A region Ig81-83 fragment gene (NM_133378.4) was codon optimized by GeneArt, and the synthetic DNA sequence was inserted in a pACpET vector (37) to produced the protein without a His-tag under ampicillin selection. The positive clones were verified by DNA sequencing and then the plasmids of titin I81-I83 and calpain-3 C129A ΔNS&IS1 were transformed into *E. coli* strain BL21(DE3) for protein expression under kanamycin and ampicillin selection.

A single clone was picked and inoculated into LB media (3 mL) containing antibiotic (0.05 mg/mL kanamycin for calpain-3 alone and 0.05 mg/mL of kanamycin plus 0.1 mg/mL of ampicillin for the calpain-3/titin complex), and incubated with shaking (220 rpm) for 8 h at 37 °C. This 3-mL culture was then added into 200 mL of LB media containing antibiotic and allowed to grow overnight at 37 °C with shaking (220 rpm). Subsequently, the overnight 200-mL culture was used to inoculate eight 1-L cultures grown under the same conditions until their OD_600_ reached 1.0, at which point protein expression was induced by the addition of isopropyl β-D-1-thiogalactopyranoside to a final concentration of 0.4 mM. After additional culturing for 16 h at 20 °C, cells were harvested by centrifugation.

### Purification of recombinant calpain-3

Calpain-3 and calpain-3/titin-I81-I83 complex were purified from lysed *E. coli* using protocols developed for calpain-2 isolation (38) with some modifications. Briefly, cells were lysed by sonication in buffer A [50 mM Tris-HCl (pH 7.6), 5.0 mM EDTA, 10 mM 2-mercaptoethanol] containing one tablet of protease inhibitor cocktail (Roche), which can inhibit a broad spectrum of serine and cysteine proteases, and the cell debris were removed through centrifugation at 48,000 x *g* for 60 min. The supernatant was applied to a DEAE ion-exchange column that had been pre-equilibrated with buffer A. After washing away unbound material, calpain-3 was eluted from the column using a gradient of 0 - 0.75 M NaCl in buffer A. Fractions containing calpain-3 based on SDS-PAGE analysis were combined and MgCl_2_ was added with gentle stirring to a final concentration of 23 mM in order to saturate the 5 mM EDTA and avoid stripping Ni*^2+^* from the Ni-NTA resin used in the next purification step. The sample was applied to a Ni-NTA column, washed with 50 mM Tris-HCl (pH 7.6), 100 mM NaCl, and 5 mM imidazole, and eluted wash buffer supplemented with 250mM imidazole.. Elution fractions containing calpain-3 according to SDS-PAGE were dialyzed into a buffer containing 20 mM Tris-HCl (pH 7.6), 100 mM NaCl, 2mM EDTA, 10 mM 2-mercaptoethanol, 0.05% Na-azide, and two tablets of protease inhibitor cocktail. Lastly, the calpain-3-enriched sample was applied to a Superdex 200 column (Amersham Biosciences) for a final purification step. Protein purity was assessed using SDS-PAGE visualized by Coomassie Blue. All purification procedures were performed at 4 °C.

### Proteolysis assay

The susceptibility of calpain-3 (C129A)ΔNS&IS1 and calpain-3/titin-I81-I83 preparations to proteolysis over time was assessed in the absence of Ca^2+^ using SDS-PAGE. Protein was assayed in 20 mM Tris-HCl (pH 7.6), 100 mM NaCl_2,_ 2 mM EDTA, and 10 mM 2-mercaptoethanol. Samples were stored at 4 °C in a final volume of 400 µL with a protein concentration of 0.5 mg/mL. Aliquots were removed at time points ranging from 0 to 70 days and the reaction was stopped by the addition of 3x SDS-PAGE loading buffer followed by heating at 95 °C. The samples were analyzed by SDS-PAGE with the protein bands visualized by Coomassie Blue staining.

### Molecular mass estimation by size-exclusion chromatography

A size-exclusion chromatography column (HiLoad 26/60 Superdex 200) was equilibrated with 20 mM Tris-HCl (pH 7.6), 100 mM NaCl, 2mM EDTA, 10 mM 2-mercatoethanol and 0.05% Na-azide at 4 °C. The column was then calibrated with molecular weight standards listed in Table S1. Following calibration, calpain-3 (C129A) ΔNS&IS1 was loaded onto the column. Protein was detected in the eluate by measuring the absorbance of fractions at 280 nm and analyzing ’peak’ fractions by SDS-PAGE.

### Multi**-**angle light scattering coupled with size**-**exclusion chromatography (SEC-MALS)

Multi-angle light scattering experiments were performed at the University of Toronto (Toronto, ON Canada). For SEC-MALS, protein samples were chromatographed on a Superdex200 increase 10/300 column coupled to a Wyatt multi-angle light scattering analyzer with dynamic light scatter (LS) and refraction index (RI) detectors in-line to an ÅKTA Pure FPLC at 4 °C. The system was pre-equilibrated using running buffer (20 mM Tris-HCl pH 7.6, 100 mM NaCl and 2 mM EDTA) overnight. A fresh solution of BSA (1 mg/mL) was prepared using the same running buffer for system calibration as well as 5 mg/mL of either C129A ΔNS& IS1 calpain-3 or calpain-3(C129A) ΔNS&IS1/titin-I81-I83). Aliquots (100 µL) were loaded onto the column. The running buffer was prepared in the absence of a reducing agent and Na-azide in order to avoid any RI mismatch signal. The column was run at a flow rate of 0.4 mL/min with a maximum pressure of 2.5 MPa, and the data were analyzed using the software package ASTRA 7.14 to obtain the molecular weight.

### Sedimentation velocity analytical ultracentrifugation

Sedimentation velocity analytical ultracentrifugation experiments were performed at the Canadian Center for Hydrodynamics (CCH) at the University of Lethbridge (Lethbridge, AB). To monitor mass action-driven reversible oligomerization, Calpain-3 (C129A) ΔNS&IS1 was measured at a high and low concentration. The high concentration (1.75 µM) was measured at 280 nm in a buffer containing 20 mM Tris-HCl (pH 7.6), 100 mM NaCl, 2 mM EDTA, 5mM TCEP, and 0.05% sodium azide. For the low concentration (0.30 µM), which was measured at 225 nm, the buffer was diluted 10-fold with ddH_2_O to reduce the absorbance contributions of TCEP and EDTA at this wavelength. Both concentrations were measured at a speed of 34,000 rpm by UV intensity detection in a Beckman Optima AUC analytical ultracentrifuge, using an An60Ti rotor and standard 2-channel epon centerpieces (Beckman-Coulter). Measurements were performed at 4 °C, and data were collected for 9 h. In addition, a low concentration sample (0.3 µM) was cross-linked by treatment with 0.1% glutaraldehyde [32] using a buffer containing 50 mM Tris-HCl buffer (pH 7.6), 25 mM NaCl, and 2 mM EDTA. The cross-linked calpain-3 sample was measured at 40,000 rpm to improve the separation of the different oligomeric states.

### Data Analysis

Data were analyzed with UltraScan-III ver. 4.0, release 7034 (39). All data were processed as described in (40). Model-independent, diffusion-corrected integral sedimentation coefficient profiles were prepared with the enhanced van Holde – Weischet analysis (vHW) (41). To determine the molecular parameters of the stable oligomeric species, the data were modeled with the parametrically constrained spectrum analysis (PCSA) (42), enhanced by Monte Carlo analysis(43). An increasing sigmoid parameterization was found to produce the best fits. The partial specific volume of Calpain-3 was assumed to be 0.729 ml/g based on its protein sequence as determined by UltraScan.

### Blue-native-PAGE (BN-PAGE) run in 1D and 2D

Calpain-3 (C129A) ΔNS&IS1 (5.8 μM) or calpain-3(C129A) ΔNSΔIS1/I81-83 (38 μM) were mixed in 4x non-denaturing loading buffer [77.5 mM Tris-HCl (pH 6.8), 25% glycerol and 0.05% bromophenol blue] and were separated on 4 -16% Native PAGE Bis-Tris gradient gels (Thermo Fisher Scientific) (44). Following electrophoresis, protein bands were visualized by Coomassie Blue staining. For two-dimensional gel electrophoresis, the first-dimension was run under native conditions on BN-PAGE, then a strip of BN-gel, which contained the complex bands, was subjected to the second dimension of SDS-PAGE electrophoresis. The protein spots on the 2D gel were visualized by Coomassie Blue staining.

### BioSAXS data collection

Biological Small Angle X-ray Solution Scattering (BioSAXS) experiments were performed on the Bio-SAXS beamline ID7A1 at Cornel MacCHESS (Ithaca, NY, USA). Calpain-3 (C129A) ΔNS&IS1 was prepared at concentrations of 1.5 mg/mL or 3.0 mg/mL, in 20 mM Tris-HCl (pH 7.6), 100 mM NaCl, 2 mM EDTA, 2% glycerol, 0.05% Na-azide and 10 mM 2-mercaptoethanol. Data were collected using an in-line Superdex 10/300 column equipped with an EIGER 4.0 detector. The beam energy range was 1.771203 Å - 0.885601 Å. A series of 1096 images and 1359 images were collected from the 1.5 mg/mL and 3.0 mg/mL samples, respectively, with 2-s exposures per image. The BioXTAS RAW software (Version 2.1.1) [34] was used for data processing and preliminary analysis. Radiation damage was not observed in the samples. A summary of the SAXS parameters is given in Table 1.

**Table 1.**
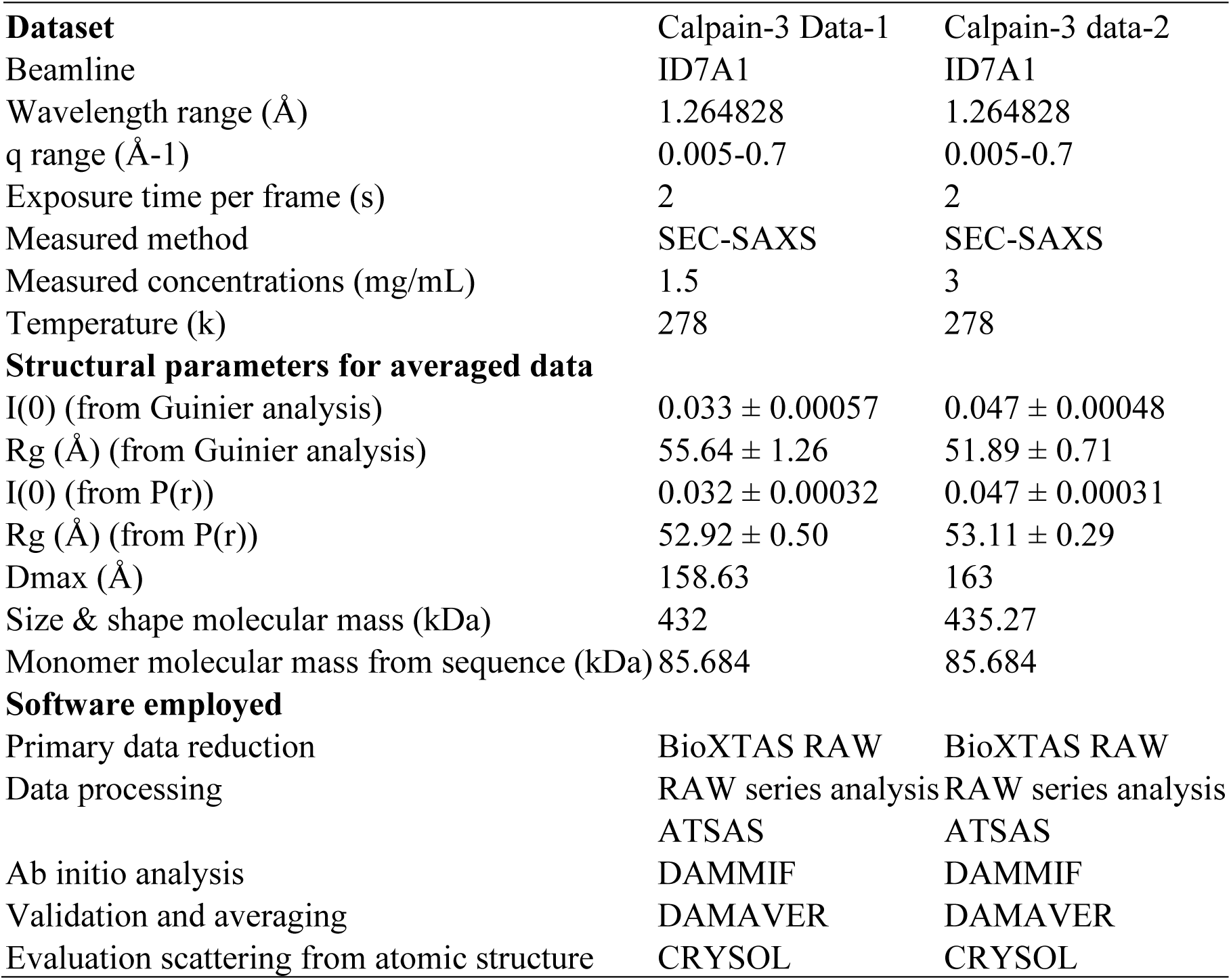

### Molecular dynamics simulations for a homodimer structural model

A predicted structure of human calpain-3 lacking the NS and IS1 regions was generated by AlphaFold2 (45). The structure was duplicated and aligned through the complementary unpaired fifth EF-hand motifs of the C-terminal PEF domains of the two calpain-3 molecules to match the crystal structure of the PEF homodimer association (PDB ID 4OKH). Subsequently, this homodimer of calpain-3 was rendered in PyMOL (The PyMOL Molecular Graphics System, Version 2.0 Schrödinger, LLC), and subjected to molecular dynamics simulations using GROMACS (<u>Version 2016.3</u>) (46). Protein was solvated in a box containing 82,423 water molecules, 192 Na^+^ ions, as well as 166 Cl^-^ ions to neutralize the charge of the system. Energy minimization was performed using the steepest descents protocol, followed by position-restrained dynamics runs to equilibrate solvent and ions around the structure by constant-volume-temperature and constant-pressure-temperature, each 0.1 ns in duration. Finally, an unrestrained dynamics simulation was run for 100,000 steps over 20 ns at a temperature of 298 K.

### Preparation of single particle cryo-electron microscopy grids and screening

Cryo-EM grid preparation and sample screening were carried out in the Biomolecular Cryo-Electron Microscopy Facility, University of California, Santa Cruz USA. A freshly eluted aliquot (4 µL) of calpain-3 (C129A) ΔNS&IS1 (2.76 mg/mL) or calpain-3-titin/I81-I83 (2mg/mL) from Superose 6 increase 10/300GL column chromatography was loaded onto glow-discharged holey carbon grids (Quantifoil 1.2/1.3 + 2 nm Carbon). The grids were blotted with filter paper for 2.5 secs at 22 °C and then immediately plunged into liquid ethane cooled by liquid nitrogen using a Vitrobot Mark IV (Thermo Fisher Scientific). A sample of calpain-3 in complex with titin-I81-I83 from a pull-down experiment was also prepared in which His-tagged titin I81-I83 was mixed as ‘bait’ with calpain-3(C129S) ΔNS&IS1 lacking its His-tag as ‘prey’ in a molar ratio of 10:1 with gentle agitation for 2.5 h at 4 °C. The mixture was then loaded onto a Ni-NTA affinity column followed by washing thoroughly with 20 mM Tris-HCl (pH 7.6), 100 mM NaCl, and 5 to 20 mM imidazole (wash-buffer) until the eluate was protein-free as judged by the lack of reaction with Bradford dye reagent (Bio-Rad). The calpain-3-titin/I81-I83 complex was eluted with 300 mM imidazole in wash-buffer. An aliquot (4 µL) of the freshly eluted complex peak solution (4.0 mg/mL) was loaded onto glow-discharged holey carbon grids (Quantifoil 2/2). The procedures for blotting and plunge freezing the grids were the same as above. Evaluation of the vitreous samples was performed on a Glacios™ Cryo Transmission Electron Microscope (Thermo Fisher Scientific) operating at 200 kV and coupled to a Gatan K2 Summit direct electron detector at cryogenic temperatures.

### Single particle cryo-EM data acquisition and image processing

Cryo-EM single particle data were collected at the Pacific Northwest Center for Cryo-EM (PNCC – Portland, USA -PNCC #160018/160310) with SerialEM (47) on a Thermo Fisher Scientific Krios G3i microscope operating at 300 kV coupled to a Gatan K3 Direct Electron Detector and BioContinuum Fringe Free energy filter. For calpain-3 data, the detector was used in super-resolution counting mode at a nominal magnification of 105,000x which produced a final pixel size of 0.413 Å. The total dose was 60 e^-^/Å^2^, over 60 frames with a nominal defocus range from -0.8 to -3.2 µm. For two datasets of calpain-3 in complex with titin-I81-I83, a nominal magnification of 165,000x was used, which resulted in a final pixel size of 0.256 Å and 0.507 Å, respectively. The total dose was 36.27 and 31.10 e^-^/Å^2^, over 50 frames, respectively with a nominal defocus range from -0.5 to -2.5 and -0.8 to -2.8 µm, respectively. The data were processed using the CryoSPARC v4.2 workflow (48). In the pre-processing, all movie frames were first aligned to obtain a motion-corrected image by Patch Motion Correction followed by estimation of the Contrast Transfer Function (CTF). Next, for calpain-3 data, a total of 4,029,390 particles were automatically selected from 4888 micrographs and then used for template-free two-dimensional (2D) classification. The best classes from 596,621 particles were subjected to 3D *ab*-*initio* reconstruction followed by 3D classification. Particles in the best class (334,733 particles) were selected for 3D refinement using *D*3 symmetry. In the post-processing, the final overall resolution FSC _0.143_ of 3.34 Å was obtained after local and non-uniform refinement.

For the datasets of calpain-3 in complex with titin-I81-I83, totals of 2,277,753 and 22,975,435 particles were automatically selected from 11,666 and 17,258 micrographs, respectively, and subjected to template-free 2D classification. The best classes were selected from 985,035 and 3,955,686 particles, respectively. A summary of the single-particle cryo-EM statistics is given in Table 2.

**Table 2.**
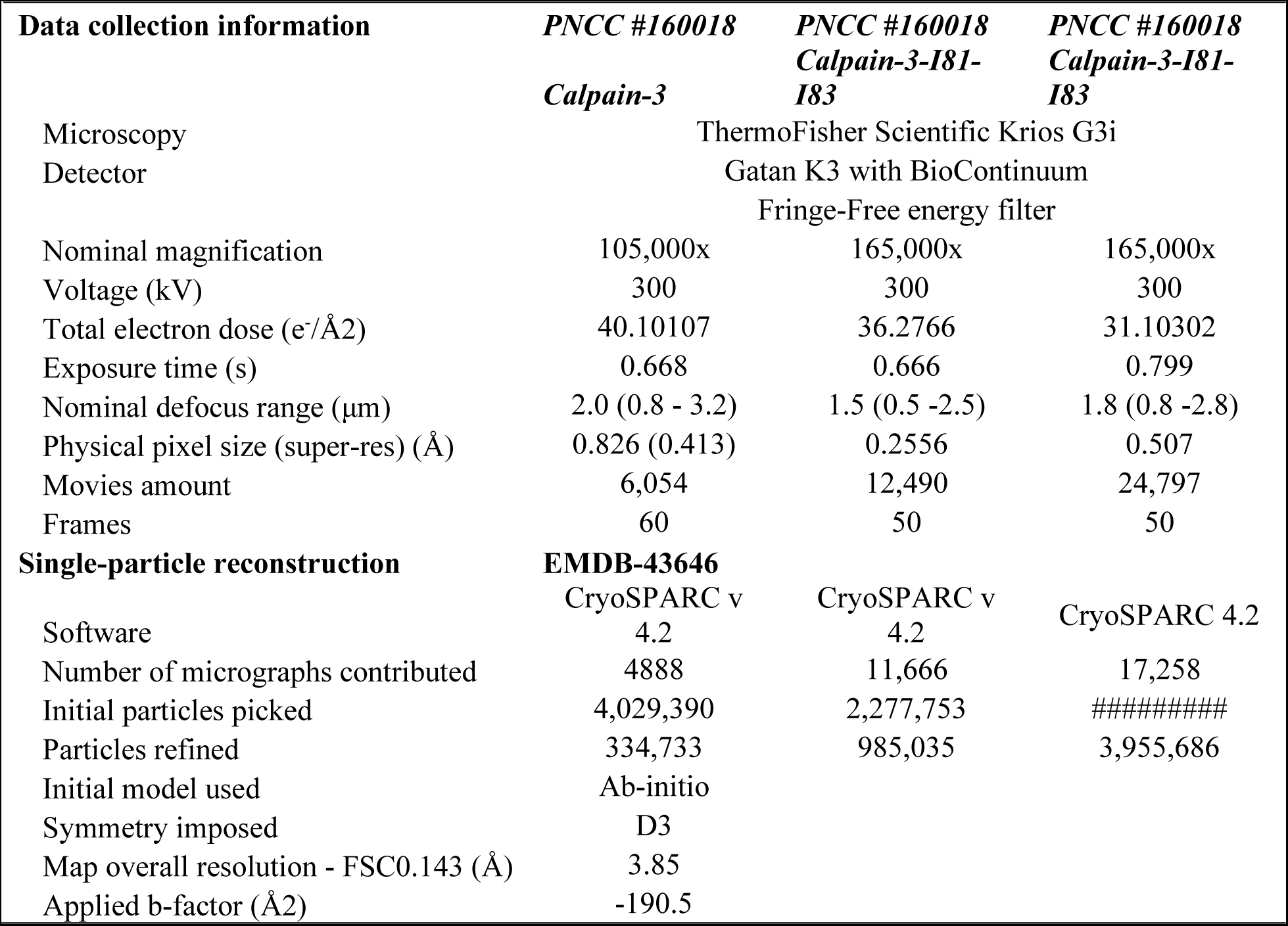
Statistics of single particle cryoEM data collection and processing.

### Model building and refinement

A sharpened cryo-EM map with a B-factor of 190.5 Å^-2^ was used for model building. A partial raw model was derived from ModelAngelo (49) as a reference, and fragments of the calpain-3 PEF domain from its crystal structure (PDB ID: 4OKH) were rigid-body docked into the map using UCSF Chimera/ChimeraX (50). Model fitting and refinement were done in Coot (51, 52).

## Results

### In vitro expression and purification of a stable recombinant human calpain-3 derivative

Characterization of endogenous calpain-3 has proven difficult because it undergoes rapid autoproteolysis during purification from muscle tissue extracts (33). One reason for this is that IS1 occupies the active site cleft and is cut as soon as the enzyme is activated (27, 28). In previous experiments, where a large quantity of recombinant calpain-3 protease core was prepared for crystallography, the presence of NS and IS1 contributed to the core’s susceptibility to both auto- and endogenous proteolysis, while mutation of the catalytic cysteine residue 129 to serine still left a trace of protease activity (26, 28). Thus, to help ensure the stable production and purification of recombinant calpain-3 from *E. coli* in this study, we designed a calpain-3 construct – calpain-3 (C129A) ΔNS&IS1 – in which NS and IS1 were deleted, and the active site Cys129 was mutated to Ala. After DEAE ion exchange chromatography, Ni-NTA affinity chromatography and size-exclusion chromatography, approximately 0.35 mg of pure recombinant calpain-3 (C129A) ΔNS&IS1 was obtained per liter of culture. The stability of this purified calpain-3 was assessed by SDS-PAGE analysis of the enzyme stored in 100 mM NaCl, 20 mM Tris-HCl (pH 7.6), 2 mM EDTA, 10 mM 2-mecaptoethanol at 4 °C for up to 70 days. The 86-kDa protein migrated as a single protein band close to the 75-kDa molecular weight standard (Figure 1). It is not known why the recombinant calpain electrophoresed ahead of where it should be, but this anomaly was consistently observed. The calpain-3 (C129A) ΔNS&IS1 derivative displayed significant stability under these Ca^2+^-free conditions, with no evidence of autolysis/proteolysis. Therefore, this storage condition was adopted in place of snap freezing and thawing aliquots.

**Figure 1.**
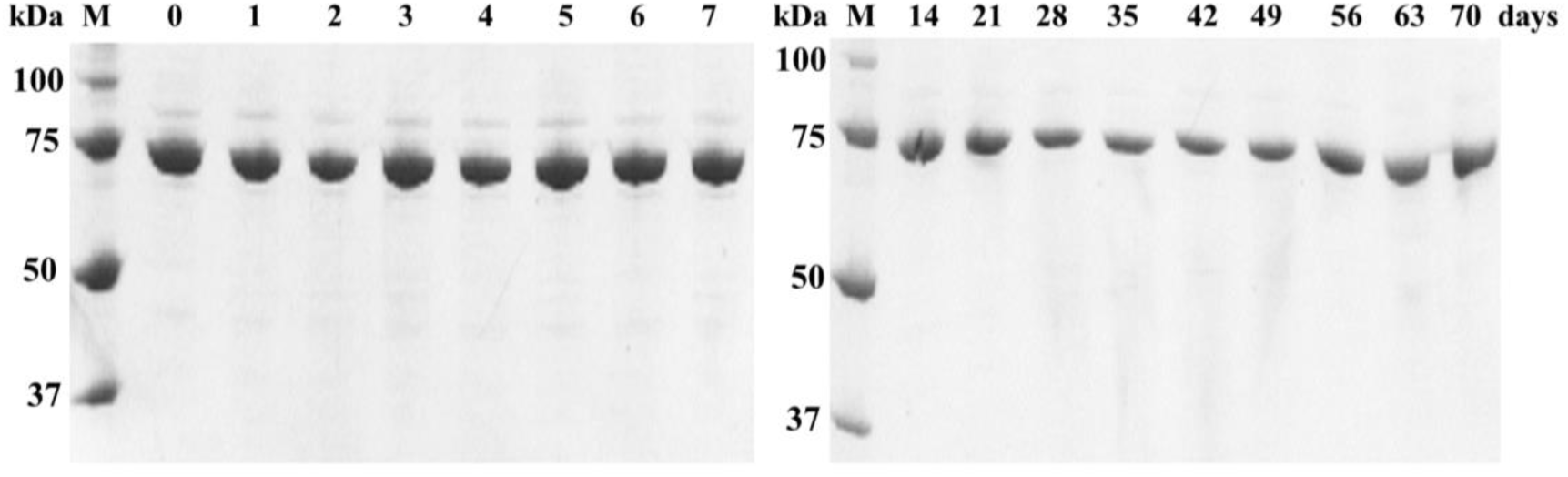
Test for autoproteolysis of calpain-3 (C129A) ΔNS&IS1 over a 70-day period. Ca^2+^-free calpain-3 was incubated at 4 °C in 20 mM Tris-HCl (pH 7.6), 100 mM NaCl_2,_ 2 mM EDTA, and 10 mM 2-mercaptoethanol. Lane numbers along the top of the two SDS-PAGE gels indicate how many days incubation each aliquot underwent. Lane M shows the separation of molecular mass markers.

### Recombinant calpain-3 is an oligomer in solution

Unlike calpains-1 and 2, calpain-3 is incapable of forming a heterodimer with the calpain small subunit (21, 33). However, the PEF domain of calpain-3 forms a stable homodimer in solution, which was confirmed by X-ray crystallography (35). Thus, the concept of homodimerization of calpain-3 was proposed and a dimer of the enzyme was modeled without any steric clashes, with an axis of symmetry running through the PEF domains, and with the two protease cores located at opposite ends of the dimer (34).

During the final step of calpain-3 (C129A) ΔNS&IS1 purification on size-exclusion chromatography in the presence of EDTA, the single peak of calpain-3 unexpectedly eluted close to where the ferritin molecular weight marker (474 kDa) chromatographed (Figure 2A). The same result was obtained when the chromatography was repeated in the presence of Ca^2+^. This apparent molecular weight is more than 2.5 times larger than that expected for a homodimer of calpain-3 (C129A) ΔNS&IS1 (theoretical homodimer molecular weight 172 kDa).

**Figure 2.**
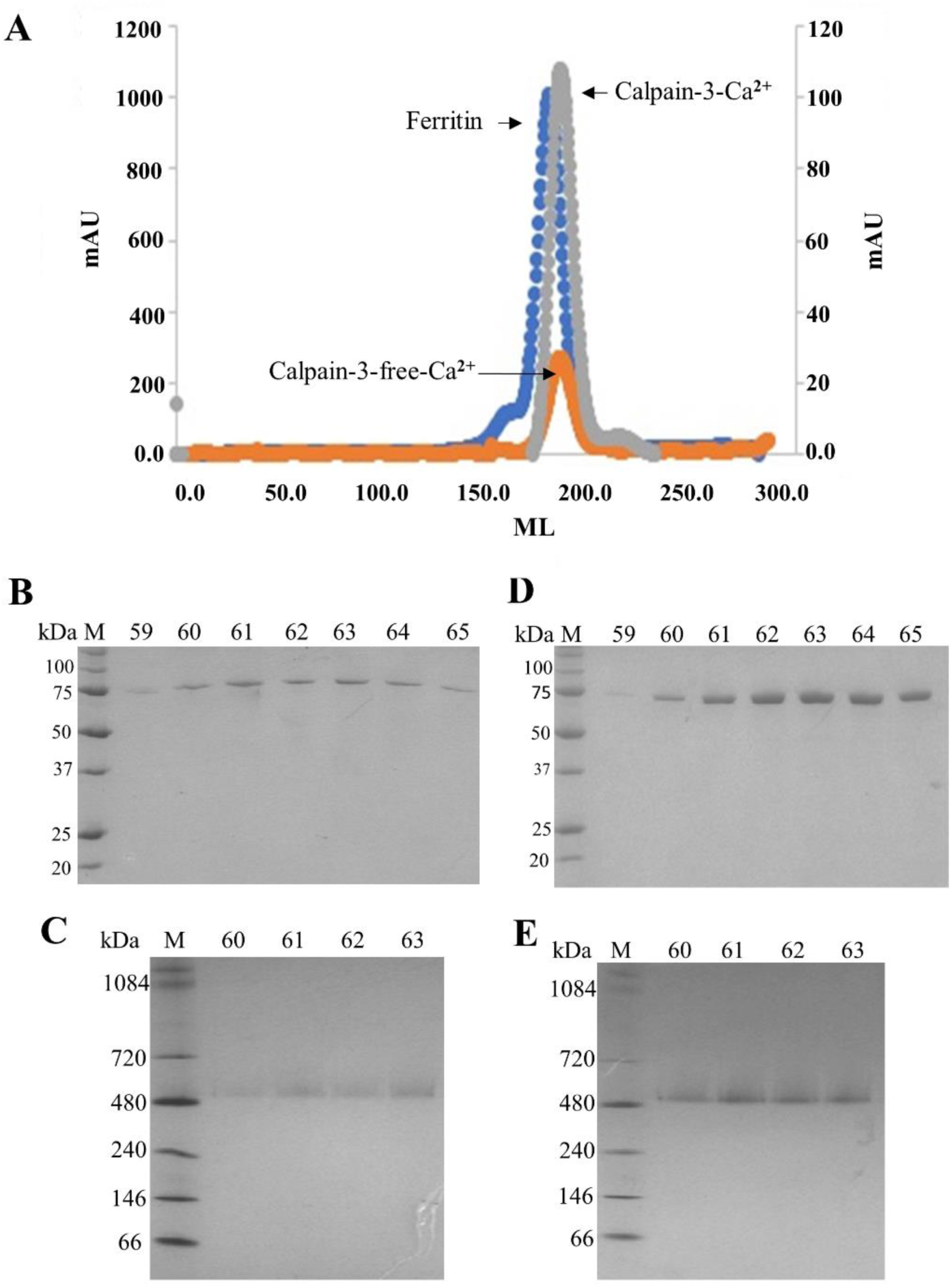
Apparent molecular weight of calpain-3 (C129A) ΔNS&IS1. **(A)** Size-exclusion chromatography of calpain-3 and molecular weight markers on a HiLoad 26/60 Superdex 200 column. Orange: Ca^2+-^free calpain-3; gray: calpain-3 in the presence of 10 mM CaCl_2_, blue: standard marker ferritin. **(B)** and **(C)** Calpain-3 peak samples in 2mM EDTA separated on SDS-PAGE and BN-PAGE, respectively. **(D)** and **(E)** Calpain-3 peak samples in 10 mM CaCl_2_ separated on SDS-PAGE and BN-PAGE, respectively. Lane numbers correspond to elution fractions from the size-exclusion chromatographies. Lane M shows the separation of molecular mass markers.

Although the model for the calpain-3 dimer suggests that it is elongated in one direction, this asymmetry seems unlikely to explain such a large increase in apparent molecular weight. SDS-PAGE analysis of the SEC peak fractions showed a single protein band of ∼ 75 kDa was present (Figure 2B). To help characterize the calpain-3 oligomer we performed Blue-native PAGE (BN-PAGE) on freshly purified, unconcentrated samples of calpain-3 (C129A) ΔNS&IS1. A single protein band was observed on the BN gels that migrated just slightly larger than the 480-kDa molecular weight marker (Figure 2C).

To confirm the molecular weight of the calpain oligomer by a method independent of its shape, the protein was assessed by multi-angle light scattering coupled with size**-**exclusion chromatography (SEC-MALS). Sodium azide and 2-mercaptoethanol were not added to the equilibration buffer for this chromatography to avoid these additive agents affecting the refraction index mismatch signal. The result indicated that 88% of the total mass fraction belonged to an oligomer of 479 (± 0.1) kDa, which was consistent with the results of BN-PAGE. However, in addition, 7.2% of the total mass corresponded to a 1478 (± 0.4) kDa species and 4.7% to a 4954.7 (± 0.3) kDa species, respectively (Figure 3A). These particles are three times and ten times larger, respectively, than the main peak material. It is possible that these smaller amounts of high molecular mass species in solution are a result of artifactual sample aggregation during the experiment and may not occur naturally.

**Figure 3.**
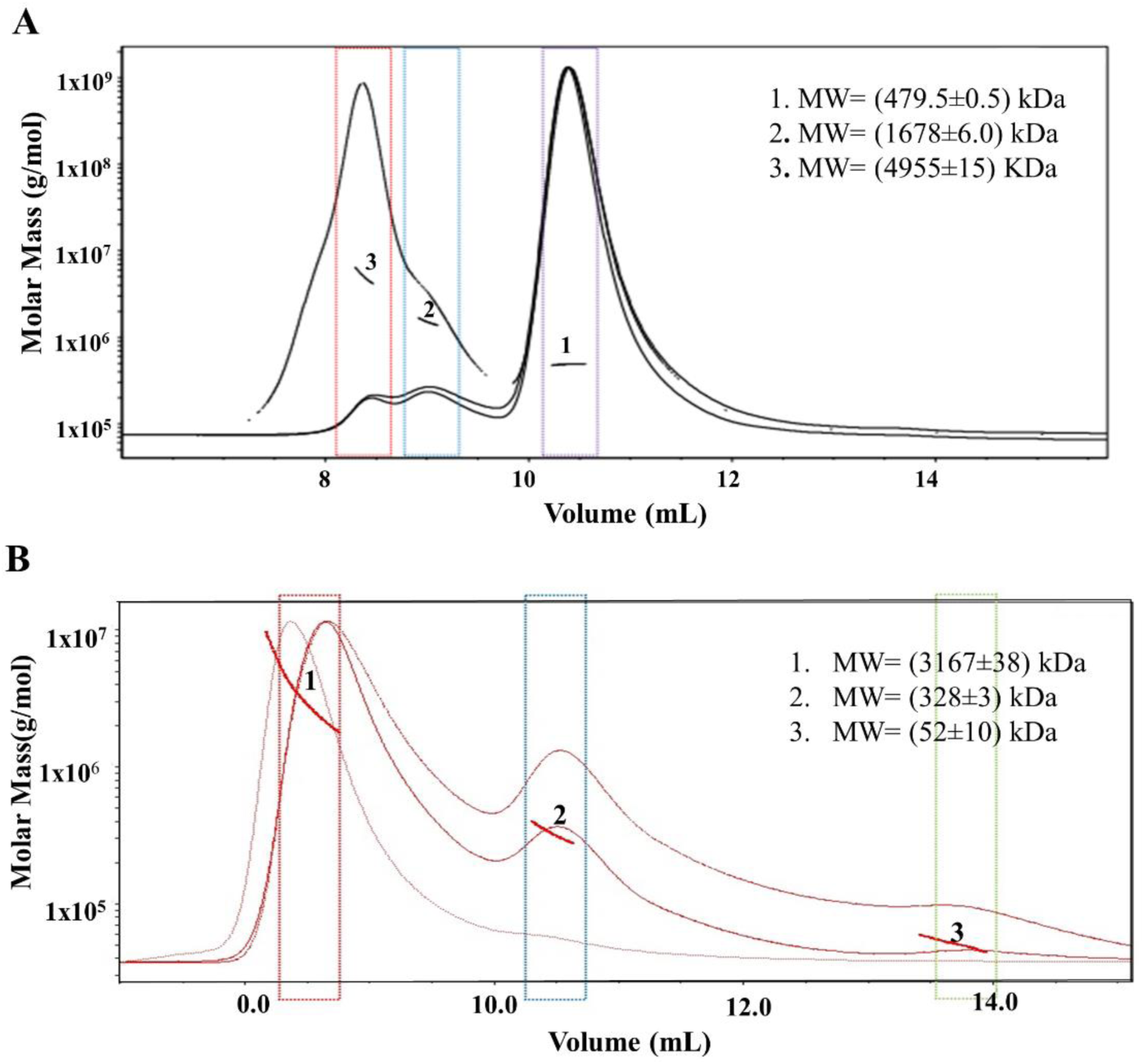
SEC-MALS analysis of calpain-3 (C129A) ΔNS&IS1 alone and in complex with titin-I81-I83. **(A)** Elution of Ca^2+^-free calpain-3 in the presence of 2 mM EDTA. Elution peaks are numbered in ascending molecular weight. Lines and arcs indicate peak molecular masses of calpain-3 in solution. **(B)** Elution of the calpain-3-titin-I81-I83 complex in the absence of Ca^2+^. Lines and arcs indicate peak molecular masses of calpain-3 and titin-I81-I83 in solution..

The results of SEC, BN-PAGE, and SEC-MALS indicate that calpain-3 (C129A) ΔNS&IS1 forms an oligomer potentially ranging from a pentamer (∼430 kDa) to a hexamer (∼516 kDa). However, since PEF domains are invariably paired up (53) it seems most likely that the oligomer is a hexamer. Because macromolecule mass in solution can be affected by many factors, such as protein concentration, shape, and buffer solutions, we explored two other biophysical methods of probing the oligomeric status of this inactive calpain-3 multimer.

### Calpain-3 sediments as an oligomer in the size range of a pentamer to a hexamer

The comparison of high and low concentration samples of calpain-3 clearly showed reversible self-association (Figure 4A, blue and green lines), as can be inferred from the right-shift of the sedimentation profiles. At low concentration (Figure 4A, blue line), about 13 % of the total concentration at 1.84 S is consistent with a monomeric species. At higher concentration (Figure 4A, green line) a stable 14 S species is formed (∼75 % of the total concentration), with a slight tendency to oligomerize further into species with values between 15-30 S. When the low concentration sample is subjected to cross-linking (Figure 4A, red line), the same 14 S species is evident, though at a lower concentration (∼45 % of the total concentration). In addition, much larger aggregates were observed as a result of cross-linking. Both the high concentration sample and the cross-linked sample were also fitted by a PCSA-Monte Carlo analysis to determine molecular parameters for the stable oligomeric form (Figure 4B, Figure 4C). The molecular parameters are listed in Table 3. The results suggest that the oligomerization of calpain-3 forms increasingly non-globular species with as mass increases. The stable 14 S species has an elevated non-globular frictional ratio of ∼1.4, suggesting a slightly non-spherical shape of the assembly. The molar masses estimated from the molecular parameters for the stable species are consistent with either a pentameric or hexameric oligomer for calpain-3.

**Figure 4.**
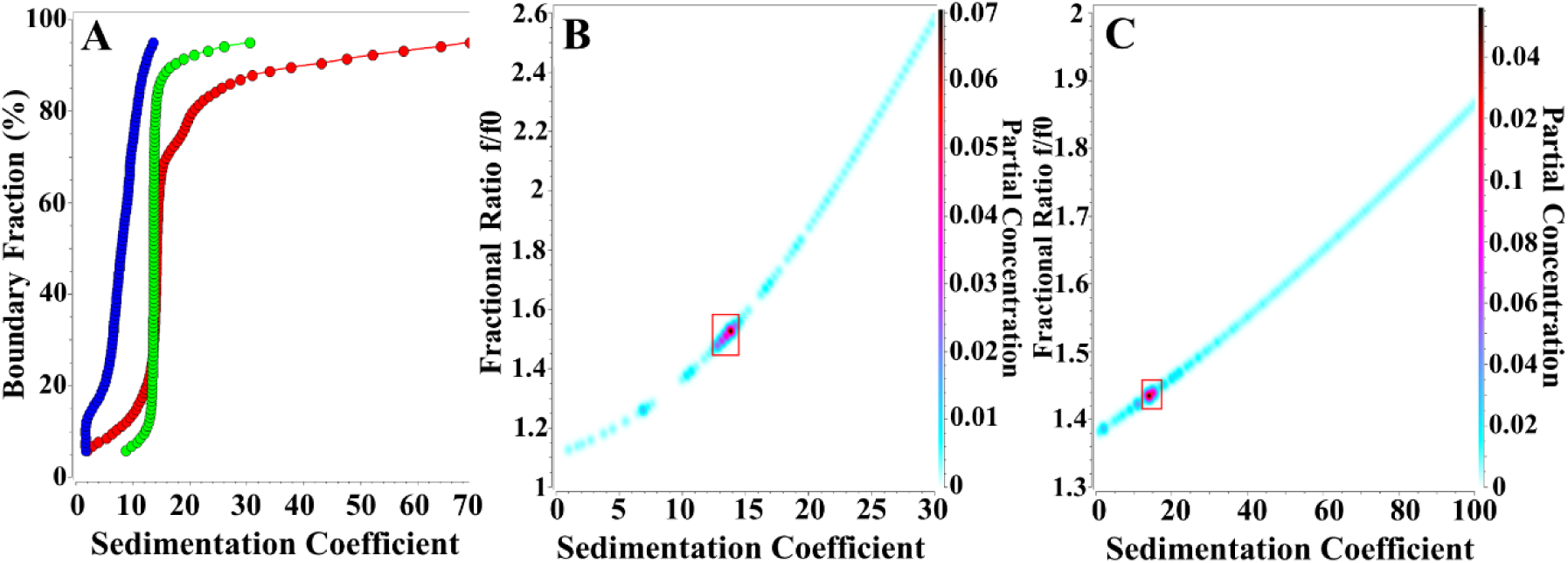
Sedimentation velocity analysis of calpain-3 (C129A) ΔNS&IS1. **(A)** Model-independent integral sedimentation coefficient distributions from an enhanced van Holde – Weischet analysis of Calpain-3 (blue: 0.3 μM, green: 1.75 μM, red: 0.3 μM, cross-linked). Reversible self-association is clearly evident from the right-shift of the green (high concentration) profile compared to the blue (low concentration) profile. A stable species around 14 S is apparent at high concentration (∼75 % of total concentration) and at low concentration (∼45 % of total concentration) when cross-linking was employed. **(B)** PCSA-Monte Carlo analysis of calpain-3 at high concentration (green line in **A**). A major species is observed around 14 S. Integration results from the indicated red square are presented in Table 3. **(C)** PCSA-Monte Carlo analysis of cross-lined calpain-3 at low concentration (red line in **A**) A major species is observed around 14 S, integration results from the indicated red square are presented in Table 3.

**Table 3.** Molecular parameters for the stable species found in Calpain-3 high concentration and cross-linked (XL) sample.

| **Sample analysis method** | **Sed. coefficient**<br>**(95% conf. interval)** | **Est. molar mass (Da)**<br>**(95% conf. interval)** | **Partial concentration** | **Fractional ratio f/f0** |
| --- | --- | --- | --- | --- |
| 1.75 μM, PCSA | 13.6 S (11.8 S, 15.3 S) | 429,100 (311,500, 546,800) | 73.60% | 1.51 |
| 0.3 μM XL, PCSA | 14.4 S (12.2 S, 16.6 S) | 429,100 (311,500, 546,800) | 43.70% | 1.44 |
| 1.75 μM, vHW | ~ 14 S |  | ~75 % | n/a |
| 0.3 μM XL, vHW | ~ 14 S |  | ~45 % | n/a |

### Solution structure analysis by small angle X-ray scattering indicates calpain-3 forms a homohexamer

That calpain-3 forms an oligomer in solution has been confirmed by the previous biophysical experiments. However, oligomeric masses showed some variation from experiment to experiment. Thus, the calpain-3 oligomer’s assembly and shape in solution may affect molecular mass determination. To probe the molecular envelope of the calpain-3 oligomer in solution, small angle X-ray light scattering (SAXS) experiments were conducted. The effect of protein concentration on the modular structure was compared by collecting SAXS data using calpain-3 samples at 1.5 mg/mL and 3.0 mg/mL. The scattering patterns of both samples were similar (Figure 5A). The scattering patterns in the low q region of the respective Guinier plots, as well as the values of Rg and I(0) that were calculated by PRIMUS (54) and GNOM (55) were in agreement for both concentrations of the sample, thus indicating no signs of aggregation or intermolecular repulsion in either sample (Table 4).

**Figure 5.**
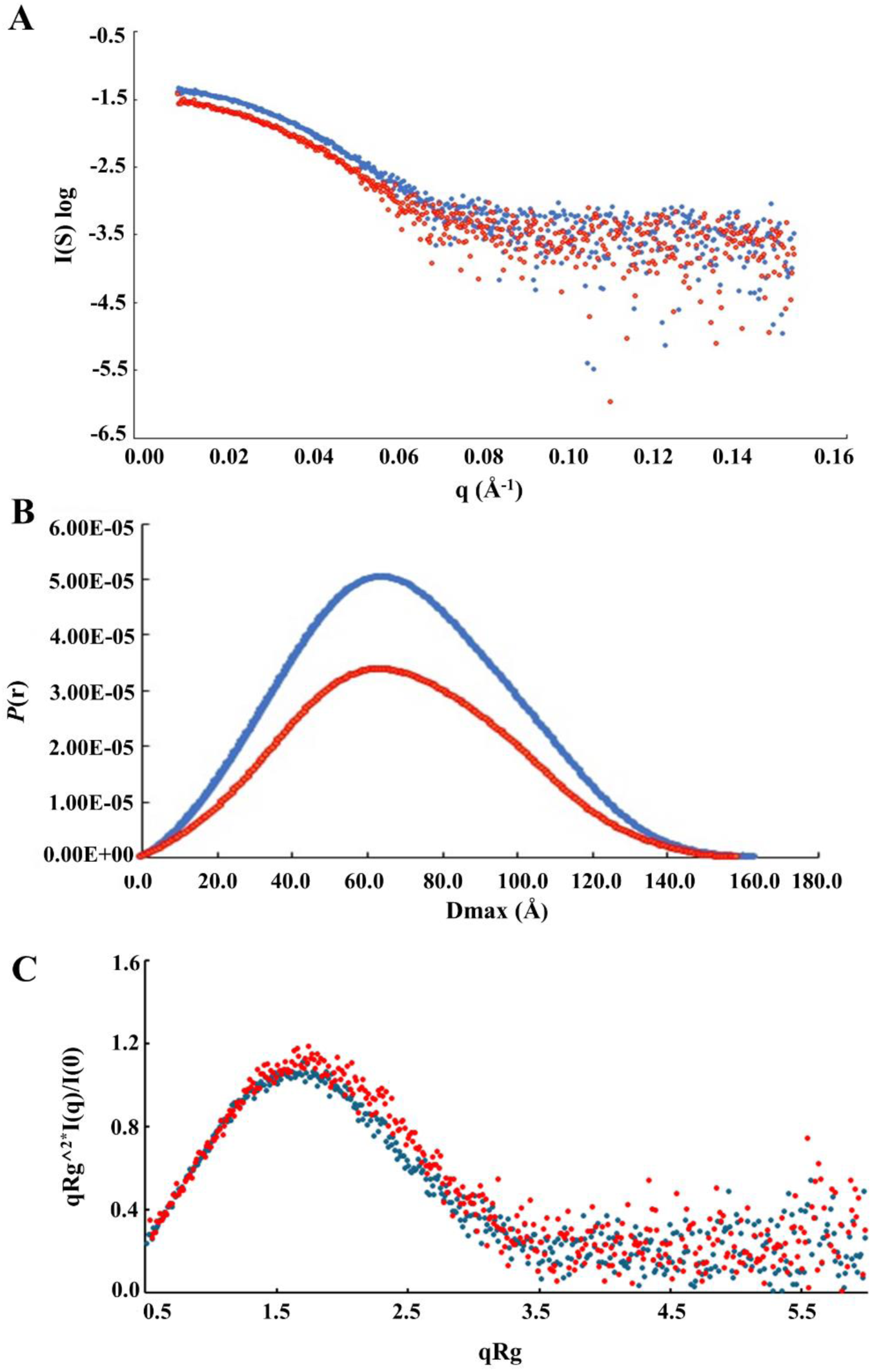
Small-angle X-ray scattering analysis of calpain-3 (C129A) ΔNS&IS1 in solution. **(A)** Superposition of scattering curves from the 3.0 mg/mL calpain sample (blue) and the 1.5 mg/mL sample (red). **(B)** Distance distribution function measurement are from 3.0 mg/mL (blue) and 1.5 mg/mL (red) samples. **(C)** Kratky-plot shows protein folding. Blue and red correspond to the 3.0 mg/mL and 1.5 mg/mL samples, respectively.

**Table 4.**

| **Sample:** | **Guinier** |  |  | **GONM** |  |  |  | **Size & shape** |
| --- | --- | --- | --- | --- | --- | --- | --- | --- |
| **Calpain 3** | **Rg** | **I(0)** | **qRg** | **Rg** | **I(0)** | **Quality (%)** | **Dmax (Å)** | **MM (kDa)** |
| 1.5 mg | 55.6 ± 1.3 | 0.03 ± 0.0006 | 1.29 | 53.0 ± 0.5 | 0.03 ± 0.0004 | 97 | 158.6 | 431.59 |
| 3.0 mg | 55.7 ± 0.8 | 0.05 ± 0.0006 | 1.3 | 54.1 ± 0.2 | 0.05 ± 0.0003 | 92 | 164.2 | 450.99 |

The pair distance distribution function *P*(*r*) was calculated with GNOM. Estimated maximum dimension (*D*max) values of 158.6 Å and 164.2 Å were obtained for the 1.5 mg/mL and 3 mg/mL samples, respectively, from the SAXS data sets. Comparison of the *P(r)* functions calculated from experimental scattering by GNOM (55) from both samples showed characteristic bell-shaped curves (Figure 5B), indicating that the protein folds into a globular shape in solution. These bell-shaped curves were identical to the bell-shaped Kratky plots (Figure 5C). Using the Size&Shape calculation of molecular mass in the ATSAS suite (56), the result shows masses of 432 kDa for the 1.5mg/mL sample and 435 kDa for the 3 mg/mL sample, respectively (Table 3). Both mass values agree with those obtained here by SEC and SV-AUC.

*Ab initio* shape reconstruction of the SAXS data was carried out in DAMMIN (57) from which a set of ten models were superimposed and averaged by DAMAVER (56). The normalized spatial discrepancy (NSD) values of the models were very similar, with mean values of 0.521 ± 0.015 and 0.534 ± 0.029, corresponding to the 1.5 mg/mL and 3 mg/mL samples, respectively. The shape of the modeled structure envelope resembled an irregular ellipse profile (Figure 6A). In order to determine the oligomer form this *ab initio* shape reconstructed model of SAXS corresponds to, we used AlphaFold2 (45) to predict a structural model of calpain-3 lacking NS and IS1 and generated a homodimer mimicking the dimer crystal structure of the calpain-3 PEF domain (PDB ID 4OKH) by aligning the two complementary unpaired fifth EF-helix motifs of PEF-hand domains from two calpain-3 structures (Figure 7). This homodimerization module again placed the two protease cores at either end of the model (Figure 7A) with their unstructured IS2 regions protruding externally (Figure 7B and C).

**Figure 6.**
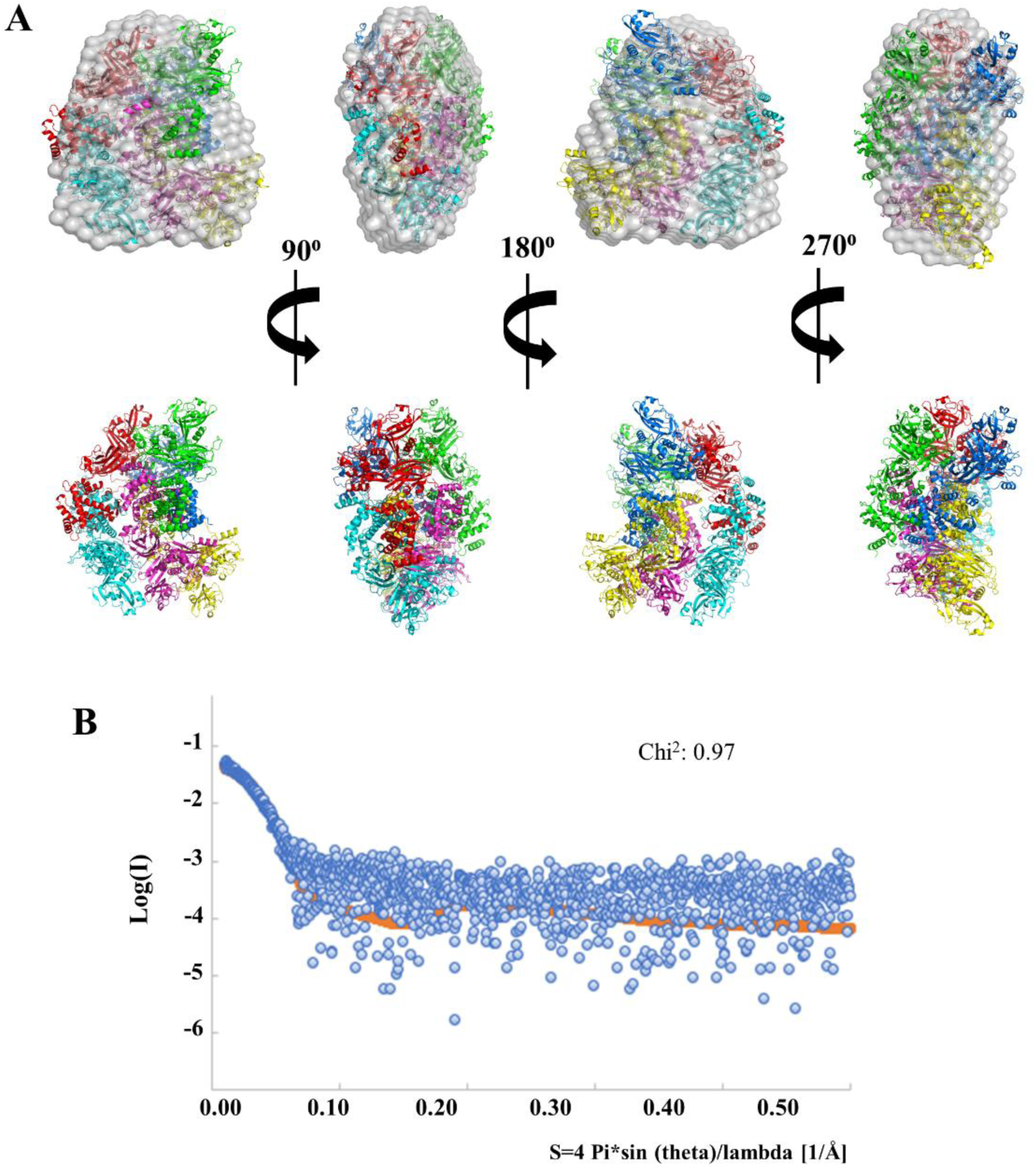
Fitting of the calpain-3 solution scattering data to the homohexamer structure model. **(A)** AlphaFold2-predicted three homodimer structures of calpain3 lacking insertion sequences are shown fitted into the *ab-initio* model envelope of calpain-3 solution experimental scattering from four different view angles. In the second row, the structures without the envelope colored red, marine blue, green, yellow, cyan and magenta represent six individual calpain-3 molecules. **(B)**. CRYSOL fitting plot shows experimental scattering of calpain-3 (blue spheres) in relations to the homohexamer structure model of calpain-3 (red line).

**Figure 7.**
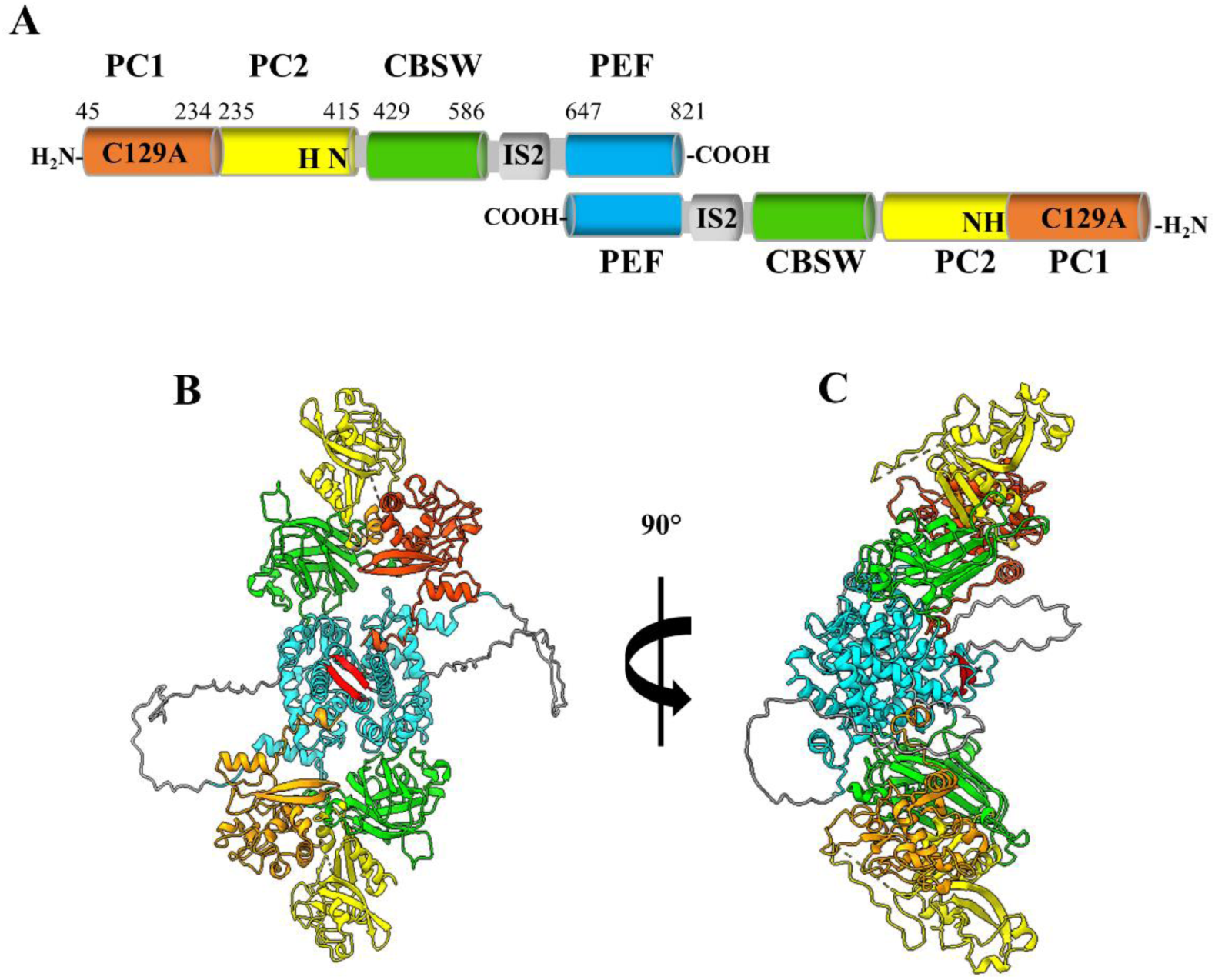
AlphaFold2 homodimer structure of calpain-3 (C129A) ΔNS&IS1. (**A)** Domain map of calpain-3 (C129A) ΔNS&IS1 homodimer paired antiparallel through the PEF domains. (**B) and (C)** Ribbon diagram of the dimer structure of calpain-3 lacking NS and IS1 but with the IS2 present viewed from the front and side, respectively. Orange and yellow ribbons represent protease core domains, PC1 and PC2, respectively. The CBSW domain is green, and the PEF domain is shown in cyan. IS2 is colored gray, and red ribbons indicate paired EF5 motifs from the two calpain-3 molecules in the dimer.

Subsequent energy minimization and molecular dynamics simulations showed no obvious steric barriers for calpain-3 homodimerization. Two homodimers of calpain-3 were fitted into the *ab initio* reconstructed envelope. Subsequently, three calpain-3 homodimers without IS2 were readily fitted into the *ab initio* reconstructed envelope. CRYSOL (58) was used to evaluate the fit of the solution scattering data from calpain-3(C129A) ΔNS&IS1 organized as three-homodimer structures. The result reveals the hexamer structural model of calpain-3 is in good agreement with the experimental scattering data and produced a χ^2^ value of 0.97 (Figure 6B).

### Three dimers of calpain-3 are present in the single particle cryo-EM map

To verify calpain-3 forms a hexamer, to obtain an atomic model, and to examine the interface connections within the oligomer, we conducted a single particle cryo-EM analysis. The data for 4,029,390 particles of calpain-3 were collected from 4888 micrographs (Figure S1A). After 2D classification, the better 2D classes from 596,621 particles (Figure S1B) were subjected to 3D *ab*-*initio* reconstruction and 3D classification. Particles in the best class (334,733 particles) were selected and used for 3D map refinement with *D*3 symmetry. As a result, the calpain-3 refined map was determined at an overall FSC_0.143_ of 3.34 Å.

The 3D reconstruction map revealed how calpain-3 forms oligomers. The middle region of the map where the six PEF domains come together is a relatively rigid region, in which the local resolution reaches 2.9 Å (Figure 8). However, the peripheral regions of the map where the six protease cores are located appear quite flexible with a much lower resolution than overall (Figure 8B and S2). The outline architecture of the calpain-3 map shows a “three-leaf clover” shape in the middle region (coloured dark blue on the resolution scale) and six angular protrusions around the “three-leaf clover” center with resolution ∼6 Å (Figure 8A and B). On closer inspection, after rendering the map at a higher density level in UCSF Chimera (50), we found a four-helix-bundle density organized as a repeated element with a total of three of these repeats present in the density (Figure 9A and B). Each four-helix bundle has an axis of symmetry suggestive of an interface between dimer pairs (coloured red and green). By fitting helices from the calpain-3 PEF domains to these densities we were able to recognize the four-helix bundle as being composed of the last two helices of the C-terminal PEF domains from a calpain-3 dimer. This confirmed the central location of the PEF domains as indicated in Figure 8A.

**Figure 8.**
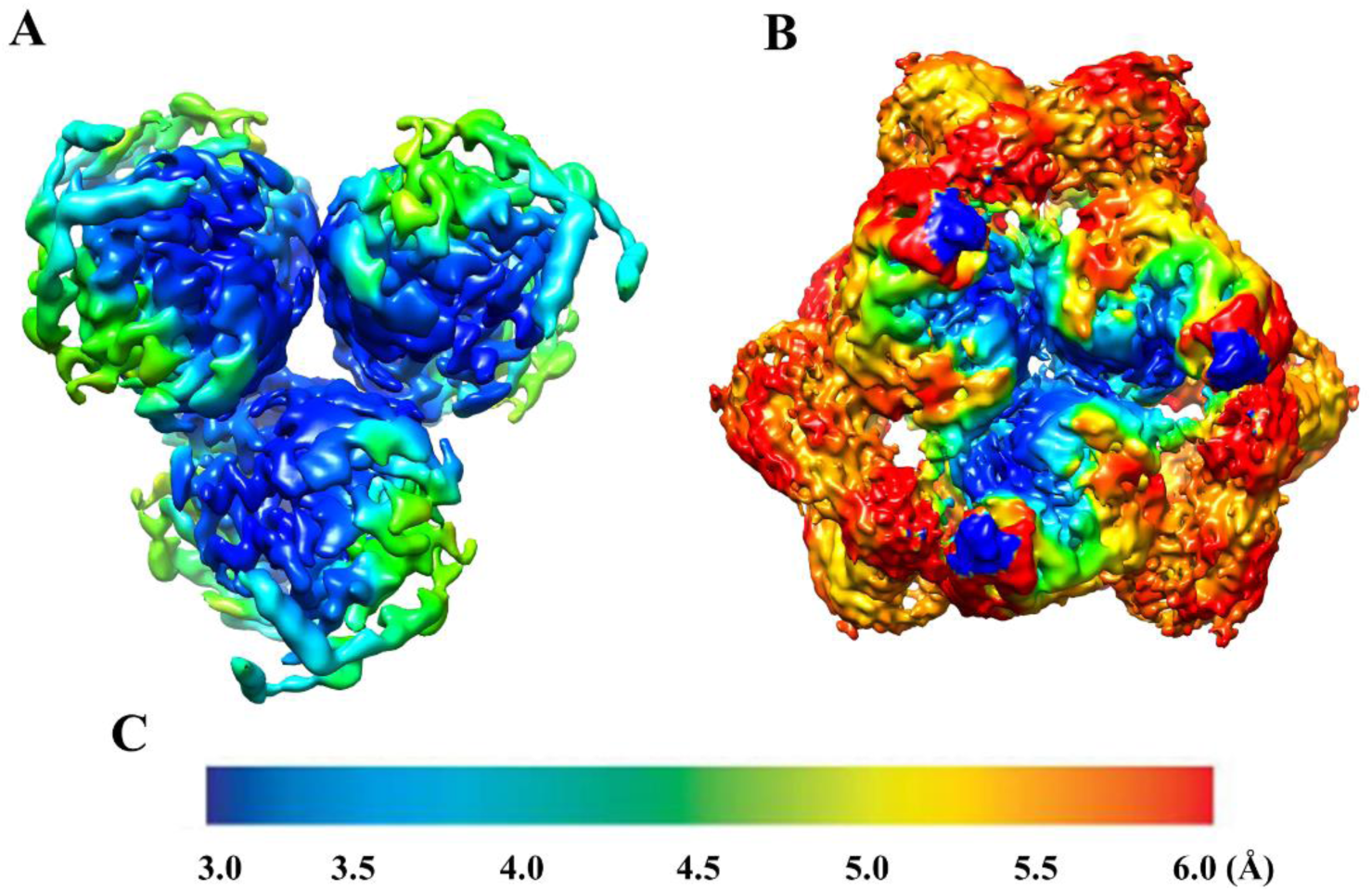
Refined cryo-EM map. **(A)** Calpain-3 segmented core map showing the 3-fold D3 symmetry of the PEF domains at contour level 0.2. **(B)** Overall calpain-3 reconstruction map showing the more flexible protease core domains on the periphery of the map. **(C)** Scale of resolution from dark blue (3 Å) to red (6 Å).

**Figure 9.**
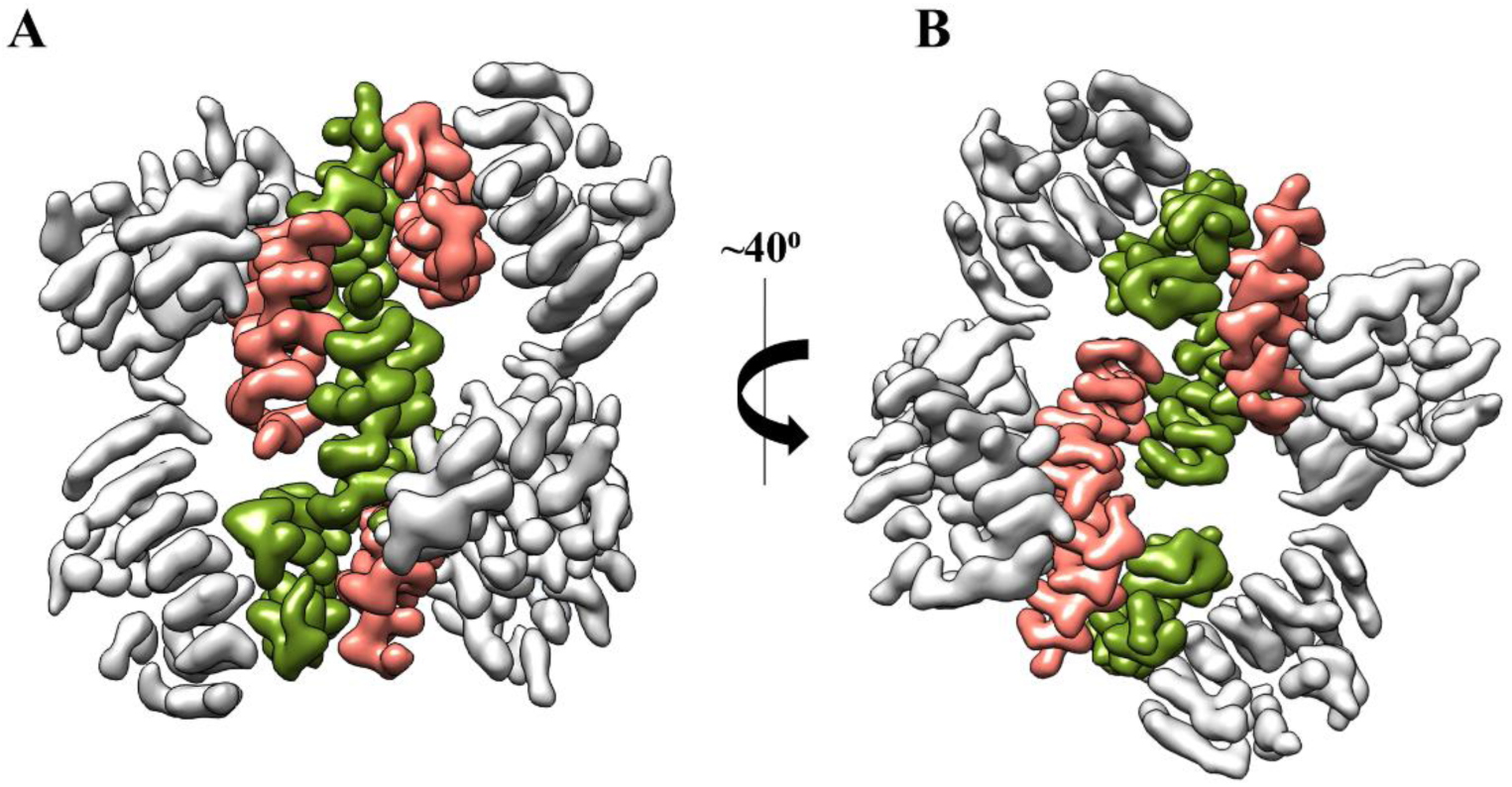
Calpain-3 PEF domain segmentation. The four-helix helical bundle in the central core is represented at a contour level of 0.67. Two views (A and B) are shown rotated by 40 ° to each other Green and orange represent the paired helices from the two different PEF domains that are the dimerization interface.

By rendering the map at a lower density level, we observed how the six subunits of calpain-3 are assembled and occupy the EM-density map (Figures 10A, B, and C). The yellow core of the oligomer seen in Figure 10A is occupied by the three PEF dimers from which emanate six calpain beta-sandwich domains (pink) connected to the protease core domains PC1 and PC2, coloured blue and green, respectively. These peripheral domains make up the six angular protrusions seen in Figure 8B coming from the core trimer of PEF dimers. When this view is horizontally rotated by 90 ° the central red colouration of yellow highlights a PEF dimer interface (Figure 10B). Two protease cores are seen on the top of the figure and two at the bottom, with the other two being hidden from view on the other side of the oligomer. The third view in this figure hints at a close association between pairs of protease core domains (Figure 10C). Although we presently do not have sufficiently high resolution in this part of the map to be confident of the details of their interaction we do have the crystal structure of the protease core to work from (28).

**Figure 10.**
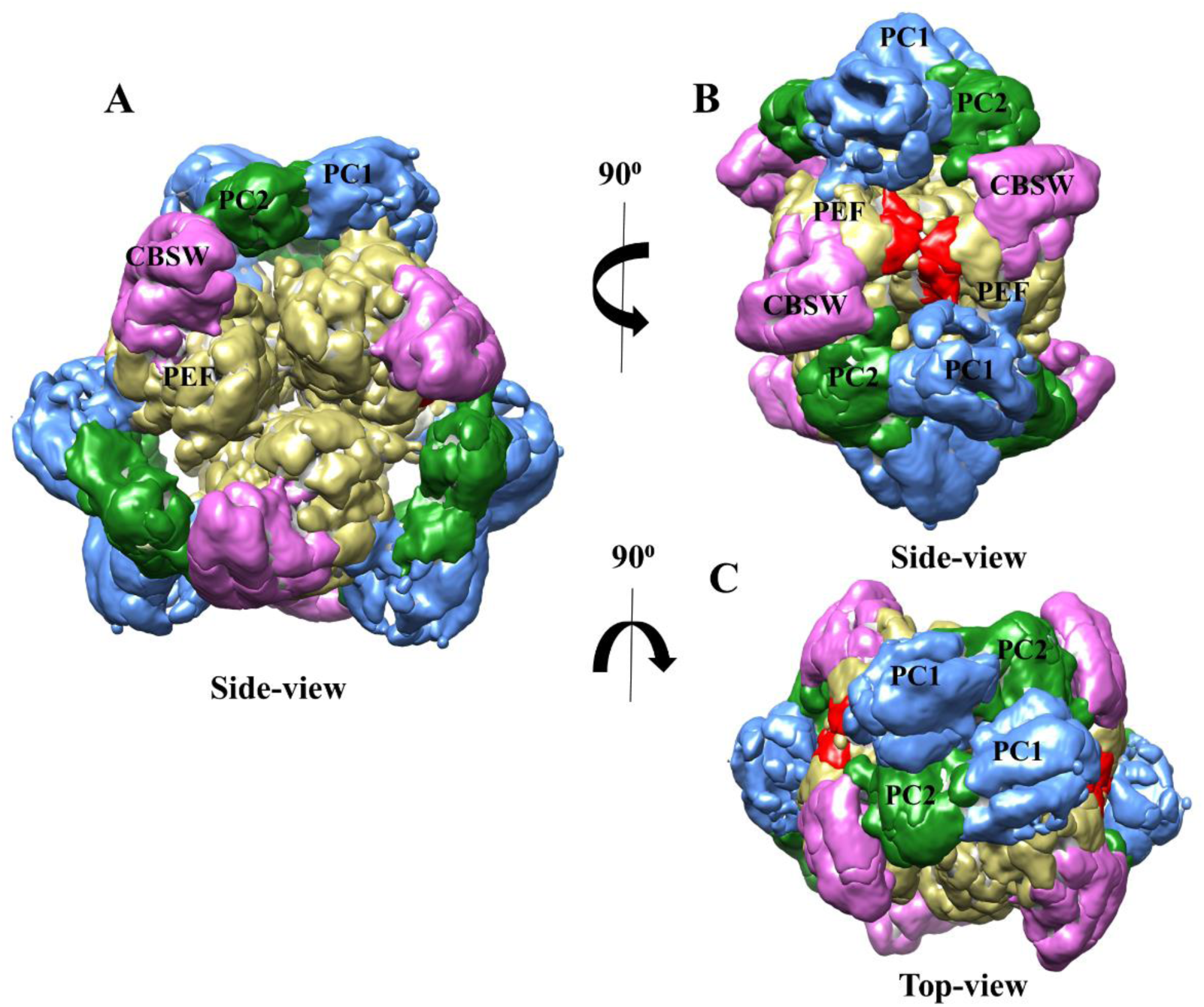
Delineation of six calpain-3 molecules in the cryo-EM density map of the hexamer showing their subdomains in different colours at a threshold level of 0.12. **(A)** Side view showing protease core domain PC1 in blue; protease core domain PC2 in green; the calpain-3 beta-sandwich domain in pink; and the PEF domain in yellow. **(B)** 90-degree vertical rotation from the view in (A) where red represents the interface between two PEF domains of a calpain-3 dimer. **(C)** 90-degree horizontal rotation from (A).

We can, however, be sure that the calpain-3 hexamer is built around the three-way association between PEF dimers at the core of the oligomer that imparts the 3-fold axis of symmetry (Figure 11A and B). Here the PEF dimer pairs are coloured green and red, grey and mauve, and blue and yellow. The three PEF dimer interfaces (Figure 11C) are centred on the four-bundle helical density elements in the middle region of the map featured in Figure 9. In the entire density map, all three of these monomer-monomer interfaces can be seen and are identical. A close-up of the homodimer interface in the cryo-EM structure reveals many hydrophobic interactions between the four contacting helices (Figure 11D) that can account for the tight binding through this interface. We also found several contacts between PEF domains that account for the ability of the PEF dimers to trimerize. These contacts come from helix4 on one PEF domain dimer and helix6 from a neighbouring dimer PEF domain along with several hydrophobic and electrostatic contacts to further help hold the three dimers together. These trimerization contacts are between the green and grey, mauve and blue, and red and yellow-coloured PEF domains (Figure 11A).

**Figure 11.**
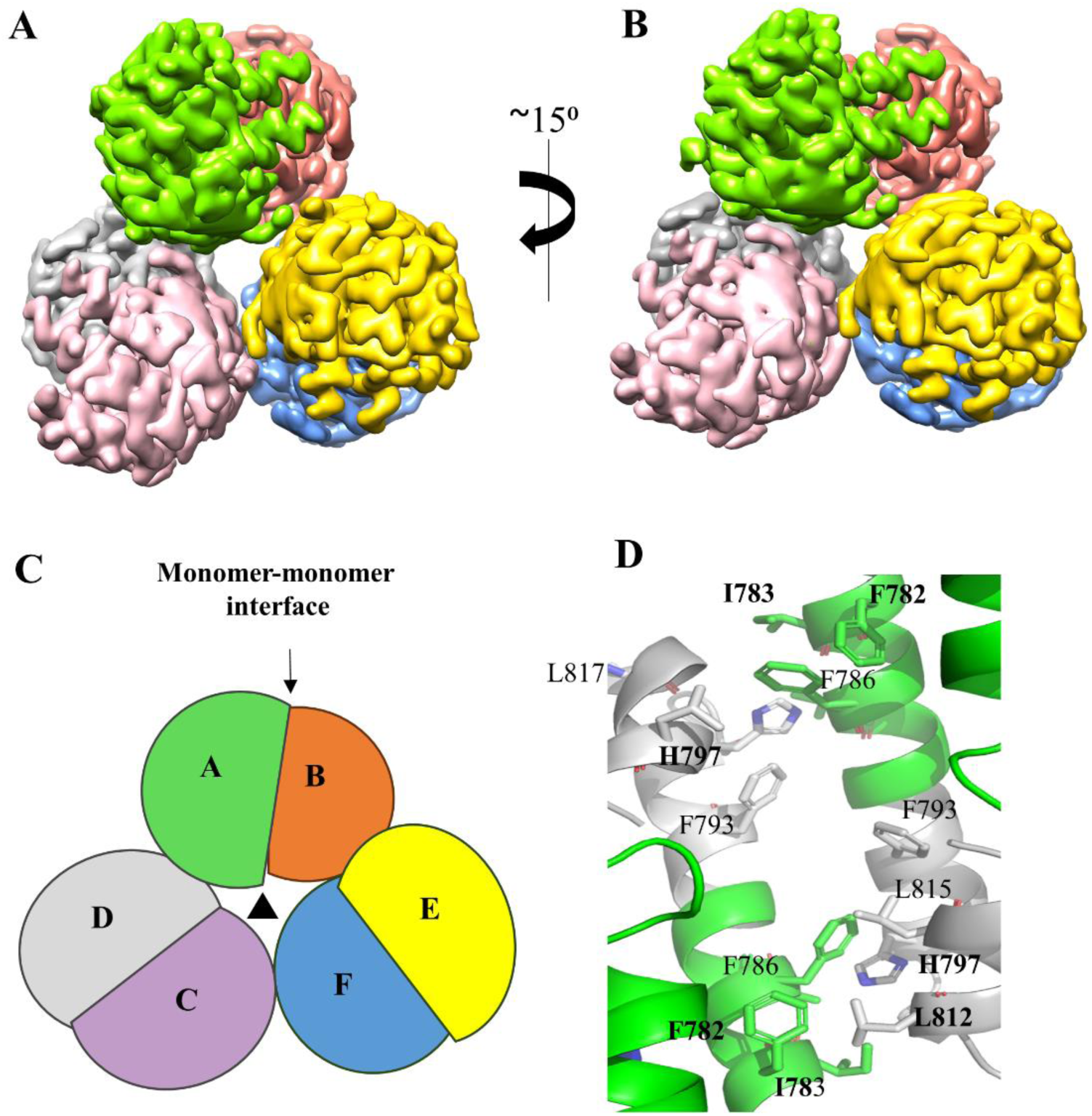
Close-up view of the six PEF domains that comprise the hexamer core. **(A)** and **(B)** Cryo-EM density map of the calpain-3 hexamer core showing its 3-fold symmetry axis at a contour level of 0.4 with two different perspectives rotated 15 ° to each other. The the PEF domains from the six subunits are shown in different colors: green, orange, purple, grey, yellow and blue**. (C)** Schematic of the monomer-monomer interfaces represented in **(A)** and **(B)**. **(D)** Close-up view of the hydrophobic residues involved in the dimerization interface between two PEF domains of calpain-3. The bold-residues represent mutated residues (https://www.dmd.nl/) that cause LGMD R1.

### PEF domain dimerization surfaces are different between solution and crystal structures

With these three interfaces mapped out, we attempted rigid body fitting of the PEF domain to this density region. We tested fitting the crystal structure of calpain-3 PEF domain (PDB ID 4OKH), calpain-2 PEF large subunit (PDB ID IDF0), which is a homologue of calpain-3 with 47.6% sequence identity, and the AlphaFold predicted calpain-3 PEF domain, but all these trials failed due to a large mismatch. Next, we used a ‘dissect-and-build’ approach from the crystal structure of calpain-3 PEF domain (PDB ID 4OKH) along with the assistance of a raw structure path in the density map that was created by ModelAngelo (49). In this way, all six PEF domains of calpain-3 were manually built into the middle density of the EM map. What is surprising and could not have been predicted is that all three PEF homodimer association interfaces are substantially different from those seen in the crystal structure of the calpain-3 PEF domain dimer (Figure 12A) (35) as well as in the calpain-2 heterodimer crystal structure (19, 20) and in the crystal structure of the small subunit PEF domain homodimer (17, 18).

**Figure 12.**
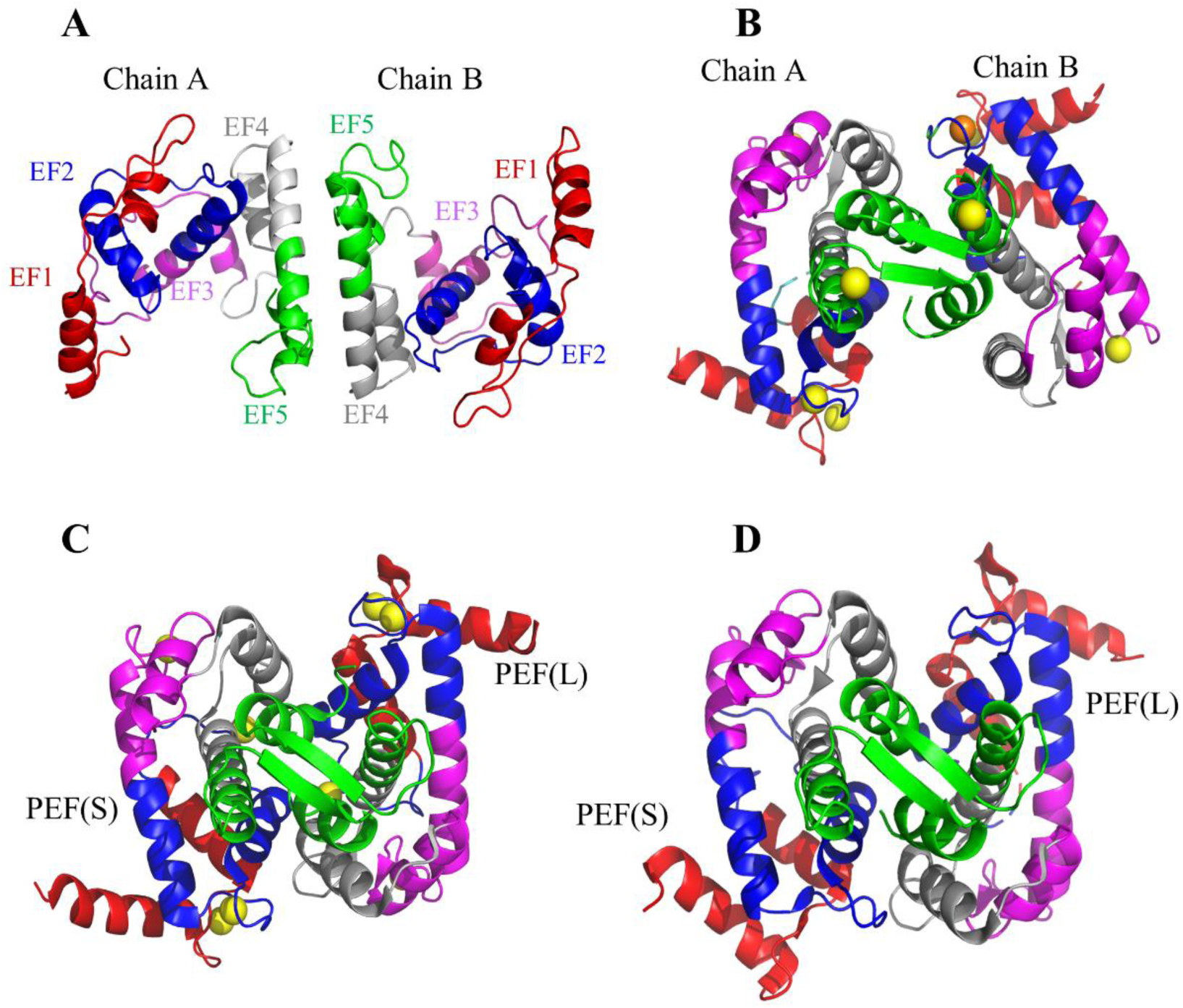
Comparison of calpain-3 PEF structures association from homodimer and calpain-2 PEF(L) and PEF(S) association from heterodimer structure in ribbon diagram format. **(A)** Calpain-3 PEF domain homodimerization solved by single particle cryo-EM; **(B)** calpain-3 PEF domain interactions revealed by the crystal structure in the presence of calcium (PDB ID 4OKH). **(C)** PEF domain interactions between calpain-2 large subunit and the small subunit with calcium present as shown by the crystal structure (3BOW); **(D)** PEF domain interactions between calpain-2 large subunit and the small subunit in the absence of calcium as shown by the crystal structure (1DF0). Structures are shown in ribbon format with different colours representing the five EF-hand helical motifs as delineated in (A).

### Binding of titin to calpain-3 splits the oligomer into dimers

Calpain-3’s association with select regions of the muscle protein titin is thought to be key to its function of muscle protein turnover after eccentric exercise (59). Major binding sites on titin for the enzyme have been identified as two distinct regions, the N2A and M-lines, respectively (30, 32). However, it is not known if calpain-3 binds there as a monomer, dimer, or oligomer. The binding site on the enzyme has been identified as the IS2 sequence that projects between the calpain beta-sandwich domain and the PEF domain (Figure 7). This region was left in the calpain-3 construct from which NS and IS1 had been removed to have the option of studying calpain-3’s binding to titin. The titin fragment alone was abundantly expressed in *E. coli* and easily purified as demonstrated for a slightly shorter version by Stronczek *et al.* (60). For those reasons, when the titin construct was co-expressed with calpain-3 it was placed in a low copy number plasmid to better balance the yields of the two products.

After purification of the recombinant products, there was a decrease in the molecular weight of the main protein band from 480 kDa to 320 kDa when electrophoresed on Blue native gel, and some unbound titin appeared to be carried along in the purification of the complex while some hexamer remained at 480 kDa (Figure 13A). To examine the composition of the 320-kDa product a lane of the Blue native gel was soaked in SDS and its proteins were analyzed in a second dimension by SDS-PAGE (Figure 13B). The two-dimensional gel showed that the 320-kDa band was composed of calpain-3 and titin in a roughly 1:2 stoichiometry to give a combined molecular weight of 2 x 86 kDa + 4 x 37 kDa = 320 kDa.

**Figure 13.**
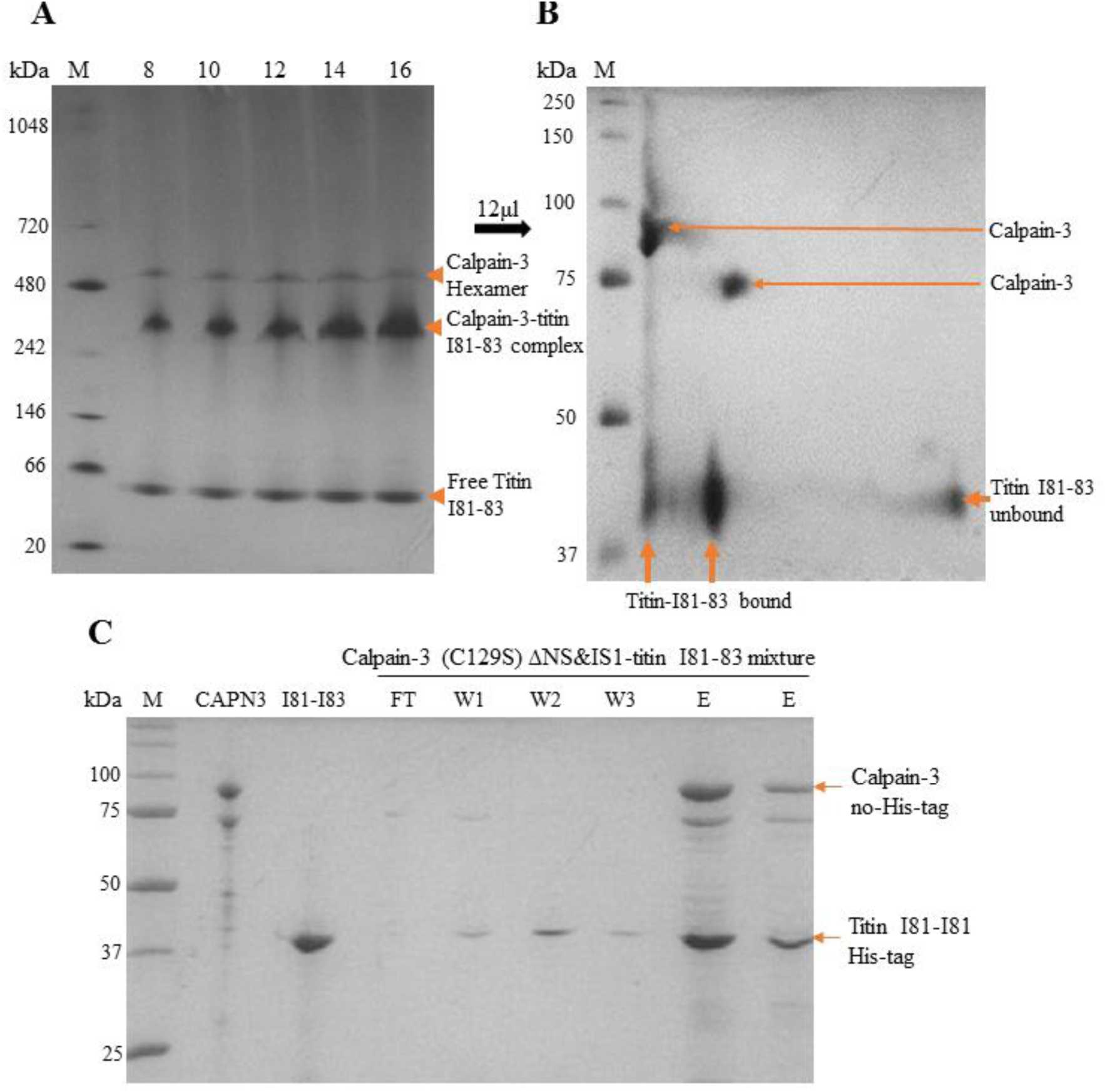
Molecular mass analysis of calpain-3-titin I81-83 complex by PAGE. **(A)** Blue-native PAGE: Lane M was loaded with the molecular mass markers with their masses indicated alongside the gel. Numbers above other lanes refer to the sample volume (µL) loaded. (**B)** Two-dimensional PAGE: A sample lane from (A) was soaked in SDS gel buffer and separated in the second dimension by SDS-PAGE. Arrows indicate the locations of calpain-3 and bound/unbound titin I81-I83 fragment. Lane M refers to the molecular mass markers for the second dimension of SDS-PAGE **(C)** Pull-down assay for complex formation between calpain-3 (C129S) ΔNS&IS1 and titin-I81-I83. LaneM: molecular mass markers. Lane CAPN3 - calpain-3; lane 181-183 - titin N2A region I81-I83 fragment. The remaining lanes are labeled as fraction from the Ni-Agarose chromatography: FT - flow-through; W1, W2, and W3 are three wash fractions. Lanes E and E are two different loadings of the eluate.

Interestingly, as calpain-3 was co-expressed and co-purified with titin N2A region I81-I83 fragment or both proteins expressed and purified individually and then incubated together, we found that they formed a complex (Figure 13B and C). However, the molecular mass of this complex was much less than that of calpain-3 alone in solution. This complex showed the molecular mass was ∼320 kDa from SEC-MALS second peak (Figure 3B) and BN-PAGE (Figure 13A). In SEC-MALS, the first peak molecular mass was 3167 (±38) kDa, which suggested the sample had some aggregation during the experiment, and the third peak molecular mass was 52(±10) kDa, which indicated some unbound free titin-I81-I83 in the co-purified sample that was consistent with two-dimension gel (Figure 13B) showing some unbound titin-I81-I83. To avoid sample degradation, we also checked the time-course experiment. The results did not show any degradation occurred during more than two months of observation (Figure S3). Based on the molecular mass calculation, the monomer of calpain-3 (C129A) ΔNS&IS1 is ∼85.7 kDa and the monomer of titin N2A region I81-I83 fragment is ∼37.5 kDa. If a monomer calpain-3 binds two titin-I81-I83 fragments, the mass ratio of calpain-3 to titin would be ∼1:2. Therefore, the doubled molecular masses at 1:2 stoichiometry (∼320 kDa) is a good agreement with the results of SEC-MALS and BN-PAGE. That implies calpain-3 dimer binds four titin-I81-I83 molecules.

### Cryo-EM images of the titin-bound calpain-3 complex are radically different from the calpain-3 oligomer

Protein purified from the titin construct co-expression with calpain-3, illustrated in Figure 13A, was analyzed by single-particle cryo-EM and revealed a mixture of particle sizes (Figure S4A). The large particles were the same as those seen with the calpain-3 hexamer alone (Figure S1A). However, the smaller particle classes were different and easily distinguished from the oligomers. To avoid any misinterpretation of the new particle classes as dissociation products of the oligomer, we devised a method to selectively purify the 320-kDa complex by elution of the His-tagged titin fragment in a pull-down experiment and directly release it onto microscope grids for plunge freezing (Figure S4B). These two-dimensional particle classes (Figure S4C and D) exactly matched those seen as the second particle type in the mixed population. The micrographs from the second dataset show the particles are homogeneous (Figure S4B) and of a size significantly smaller than that of calpain-3 hexamer (Figure S1A). After several rounds of 2D classification, the 985,035 particles in dataset 1 and 3,955.686 particles in dataset 2 were subjected to a final 2D class selection. The 2D-classes show particle-shapes are similar in both datasets of the complex (Figure S4C and D). The particle shapes in both sets are quite different from the calpain-3 hexamer (Figure S1B), which has defined shapes with symmetry (Figure 11A). The particles of the calpain-3/titin-I81-I83 complex are smaller, more extended and less symmetrical than those of the calpain-3 hexamer.

## Discussion

PEF domain proteins are always found in pairs (53) with the fifth EF-hand motif serving as the dimerization interface (61). As described above, this observation has been borne out in the human calpain family where about half of the members possess a PEF domain at their C-terminus. Calpain crystal structures have documented examples of both homo- and heterodimerization through PEF domains. For calpain-3, its PEF domain expressed as a recombinant protein in *E. coli* forms a strong dimer by pairing of the 5^th^ EF-hand (35). During the characterization of the recombinant calpain-3 oligomer by the various biophysical techniques used here the primacy of the PEF pairing led us to discount suggestions that the oligomer might be a pentamer or a heptamer. Although the weight of evidence was in favour of the oligomer being a hexamer it was reassuring to see this unequivocally established by the cryo-EM structure that showed the calpain-3 complex was a trimer of dimers.

During the early stages of this study Hata *et al*. (36) reported that the 94-kDa calpain-3 (C129S) recombinantly expressed in COS7 cells or extracted from skeletal muscle of knock-in mice had mobility on Blue Native PAGE consistent with a mass greater than 240 kDa that most closely resembles a trimer theoretical molecular weight of 282 kDa. Cross-linking with glutaraldehyde prior to biophysical analysis gave similar results suggesting that this trimer-sized particle was not a breakdown product of a larger assembly. These authors reported that the calpain-3 PEF domain was needed for trimer formation but when it was expressed as an isolated domain it formed a homodimer, consistent with our observations (35). They also showed that the removal of the insertion sequences IS1 and IS2 from calpain-3 still produced a trimer. The most puzzling aspect of these results is the suggestion that a PEF domain remains unpaired. This conundrum could likely be resolved by the use of single particle cryo-EM given that the complex is large enough at ∼280 kDa for this methodology to be used and for the results to be compared with those from this study.

Ironically and surprisingly, after relying heavily upon the 5^th^ EF-hand pairing to argue for a fundamental underlying dimer, the cryo-EM structure shows that this is not the motif pairing that dimerizes the PEF domains within the oligomer. Instead, it is the formation of a four-helix-bundle structure involving helices from EF-hands 4 and 5 that joins the calpain-3 dimers together in all three instances in the hexamer. In the early days of calpain structural biology, an attempt was made to render the calpain small subunit PEF domain monomeric by removing its 5^th^ EF-hand (62). After this deletion, an asymmetrical dimer formed in place of two monomers indicating that this domain has more than one surface available for protein-protein interactions. It would be interesting to know which of the two PEF interactions is the lower energy state. In our working hypothesis for the role of the calpain-3 hexamer in functioning human muscle we envisage oligomerization as a way of sequestering an autoproteolytically active enzyme from the point of synthesis on polysomes to its deposition into the N2A region of aligned titin molecules in the sarcomere (Figure 14). When the hexamer dissociates there, do calpain-3 dimers form through the 5^th^ EF-hand pairing at a lower energy state to help drive this reorganization? This could occur in conjunction with a positive association from the IS2 region of calpan-3 binding to titin. If so, preventing a preferred dimer association through the 5^th^ EF-hands from forming too early might be a way of spring-loading the oligomer to dissociate under the right conditions provided by titin in the sarcomere.

**Figure 14.**
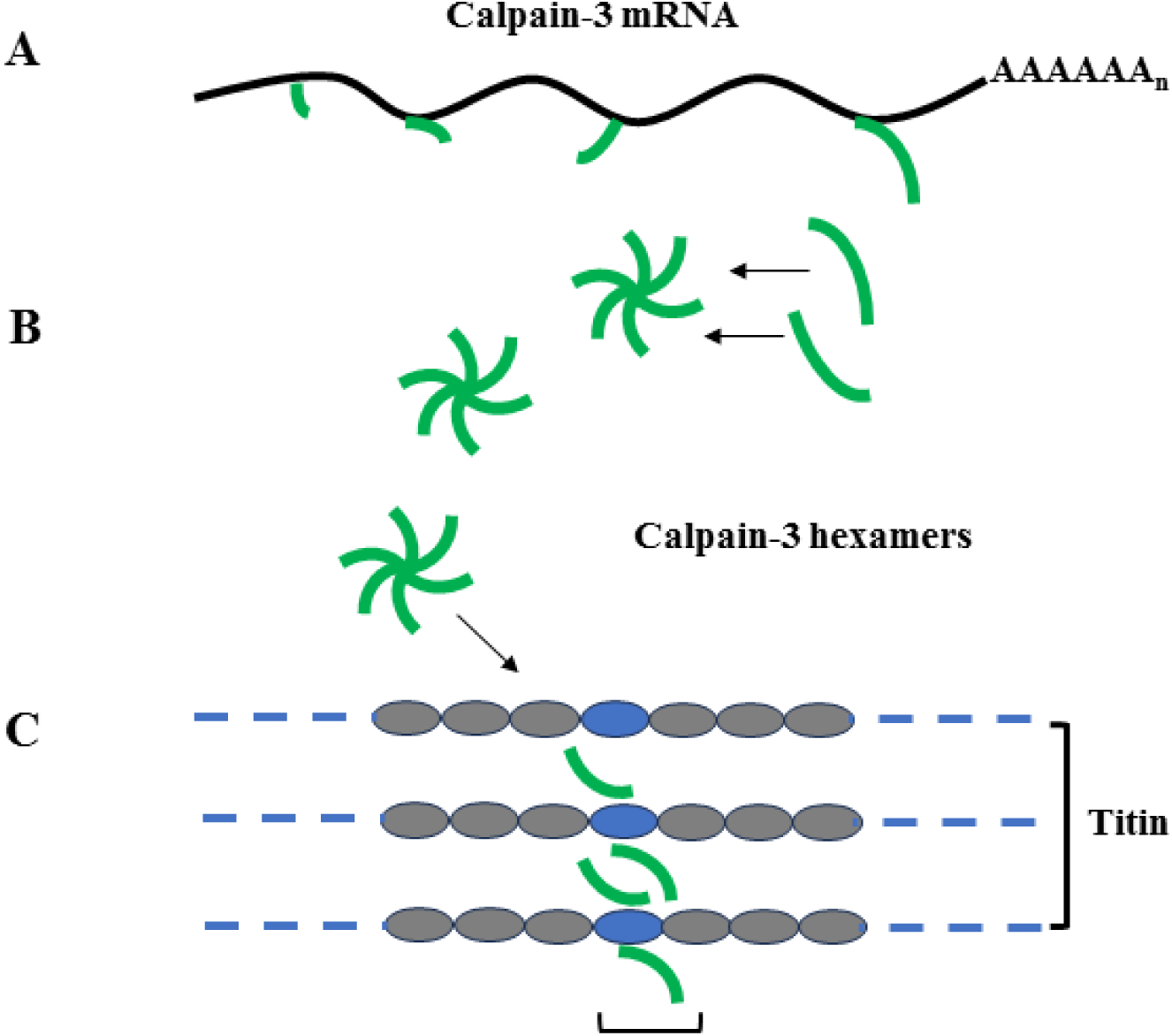
Schematic model for calpain-3 deposition on titin. **(A)** Calpain-3 (green line) is produced as a monomer by translation of its mRNA (black line with poly(A) tail) where its high local concentration favours oligomerization into a hexamer. **(B)** Calpain-3 hexamers transit to the sarcomere. **(C)** On binding to the N2A region of the vertically aligned titin (indicated by the blue oval) the calpain-3 hexamer dissociates into dimers that bind to the high local concentration of the binding sites.

Beyond the dimerization issue is the question of how the dimers come together to form the hexamer. Within the core of the hexamer the three PEF-dimers pack together through surface associations of helices 4 and 6. Each dimer pair projects in the opposite direction and the role of the beta-sandwich domain seems confined to being the linkage to the protease core region.

Looking at the protease core regions on the outer surface of the hexamer there is an intimate pairing of PC1 and PC2 from one core with the PC1 and PC2 domains from a neighbouring core that belongs to a different dimer in the hexamer (Figure 10). This association has two axes of symmetry, one running through the PC1 pair and one through the PC2 pair. Unfortunately, these three examples of core dimerization lie on the outer edge of the oligomer where the resolution is insufficient to delineate specific residues in binding. Crosslinking studies could potentially be used to confirm the juxtaposition of neighboring residues and their distance apart in the oligomer.

In designing the more proteolytically resistant calpain-3 (C129A) ΔNS&IS1 for the oligomerization study the intrinsically disordered IS2 region was left in the construct at the risk of increasing proteolysis because it was expected to be the landing pad for titin. Co-expression studies with the select titin fragment have validated this strategy by showing that the calpain-3 oligomer undergoes a remarkable breakdown into titin-bound dimers. Finding out the mechanism behind this transition is an important future direction of inquiry. However, the flexibility of the titin-bound calpain-3 dimer does not lend itself well to cryo-EM single particle analysis. Nevertheless, crystallography of select regions of the dimer complex with titin might reveal, for example, if the PEF dimerization has switched to the 5^th^ EF-hand pairing during the dissociation.

Another pressing area for further examination is the mapping of surface LGMD R1 point mutations onto the hexamer to see if any of them might destabilize the oligomer. Previously, we would have focussed on the 5^th^ EF-hand region. But now it is clear that the dimerization region in the oligomer includes nearby helices and that the PEF trimerization interface includes other PEF domain surfaces. At the other end of the oligomer, it will be necessary to interpret surface mutations in the protease core domains to see if any might make contacts between the cores. Not only should these mutations be mapped but they should be reproduced in the calpain-3 (C129A) ΔNS&IS1 construct to see if they weaken or block the assembly of the hexamer.

## Supporting information

Suppplementary figures and tables

## Acknowledgements

We dedicate this manuscript to the memory of the late Professor Hiroyuki Sorimachi whose pioneering work on p94 (calpain-3) paved the way for this investigation.

The Canadian Center of Hydrodynamics (CCH) at the University of Lethbridge was instrumental in performing sedimentation velocity analytic ultracentrifugation and subsequent analysis of calpain-3 samples with funding to BD from the Canada 150 Research Chairs Program (C150-2017-00015), Canada Foundation of Innovation (CFI-37589), Canadian Natural Science and Engineering Research Council (DG-RGPIN-2019-05637), NIH-NIGMS (1R01GM120600), NSF (TG-MCB070039N) and the University of Texas (TG457201).

We thank Dr. Omar Davulcu as well as Rose M. Haynes and the Pacific Northwest Center for CryoEM (PNCC) at Oregon Health & Science University for data collection and support on data processing. A portion of this research was supported by NIH grant U24GM129547 and performed at the PNCC at OHSU and accessed through EMSL (grid.436923.9), a DOE Office of Science User Facility sponsored by the Office of Biological and Environmental Research. We also thank Drs. Irina Novikova and Craig Yoshioka at PNCC for data processing training and support at PNCC Compute. The authors also acknowledge the Biomolecular cryo-Electron Microscopy Facility at the Department of Chemistry and Biochemistry of the University of California - Santa Cruz (RRID:SCR_021755) for scientific and technical assistance (NIH High-End Instrumentation program, S10OD02509). Molecular graphics and analyses performed with UCSF ChimeraX, developed by the Resource for Biocomputing, Visualization, and Informatics at the University of California, San Francisco, with support from National Institutes of Health R01-GM129325 and the Office of Cyber Infrastructure and Computational Biology, National Institute of Allergy and Infectious Diseases. This work was advanced by research conducted at the Center for High-Energy X-ray Sciences (CHEXS), which is supported by the National Science Foundation (BIO, ENG and MPS Directorates) under award DMR-1829070., and the Macromolecular Diffraction at CHESS (MacCHESS) facility, which is supported by award 1-P30-GM124166-01A1 from the National Institute of General Medical Sciences, National Institutes of Health, and by New York State’s Empire State Development Corporation (NYSTAR). We are grateful to Dr. Jeffery Lee for access to his SEC-MALS apparatus at the University of Toronto. Operational support was funded by a grant from the US Muscular Dystrophy Association (MDA577340) and by Foundation Award FRN 148422 from the Canadian Institutes of Health to P.L.D., who holds the Canada Research Chair in Protein Engineering. We thank our colleagues Dr. Mona Rahman, Dr. Yasuko Ono, Mathias Bell and Thomas Hansen for their comments and edits on the manuscript.

---

Figure S1: Single-particle cryo-EM analysis. **(A)** representative micrograph. **(B)** Selected 2D-classes used for the *ab-initio* reconstruction.

Figure S2: Helical bundle resolution within paired PEF domains. Calpain-3 segmented core map showing the local resolution of the helical bundle in the middle regions at contour level 0.45.

Figure S3. Time-course analysis of calpain-3 (C129A) ΔNS&IS1-titin-I81-I83 complex proteolysis. Lane M was loaded with SDS-PAGE molecular mass markers. The time at which each aliquoted protein was taken during the time-course is indicated in days by lane numbers along the top of the gels.

Figure S4. Single particle cryo-EM micrograph images of calpain-3 (C129A) ΔNS&IS1 in complex with titin I81-I83. **(A)** Sample micrograph was from the co-expression and co-purification of calpain-3-titin I81-I83 complex. Scale bar in the right bottom corner indicates 50 nm. **(B)** Sample micrograph was from the pull-down experiment that excluded calpain-3 homohexamer. Red arrows point to complex particles of calpain-3 with titin I81-I83, and yellow arrows indicate calpain-3 hexamer particles. **(C)** and **(D):** selected two-dimensional class images of the complex from co-expression and the pull-down procedure, respectively.

