## Supplementary material for "Human calpain-3 and its structural plasticity: dissociation of a homohexamer into dimers on binding titin": Suppplementary figures and tables

**SUPPLEMENTARY FIGURES**

**Figure S1.**

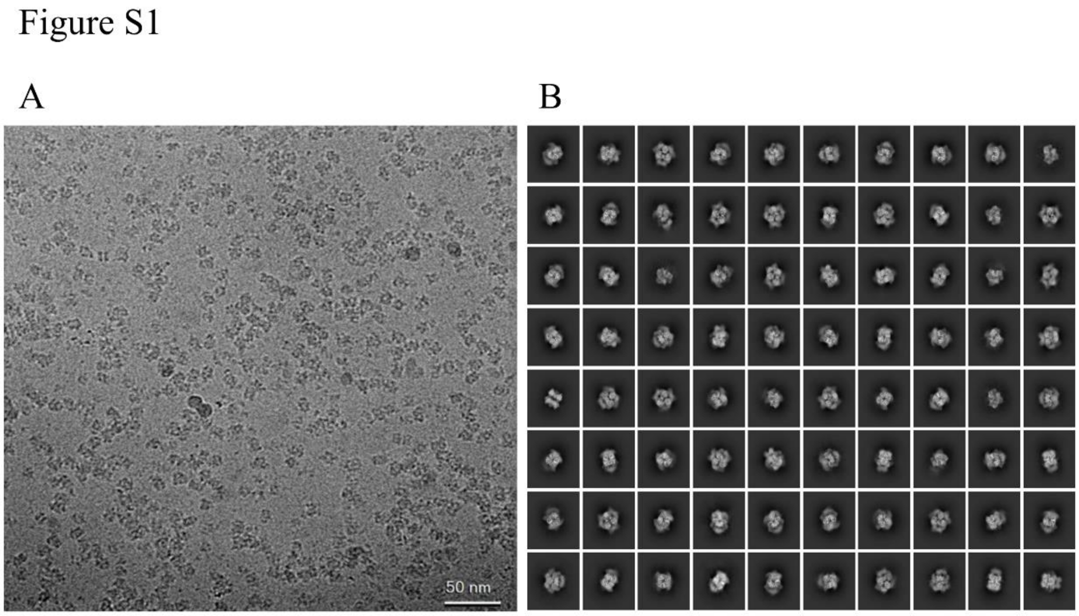

**Figure S2.**

**A**

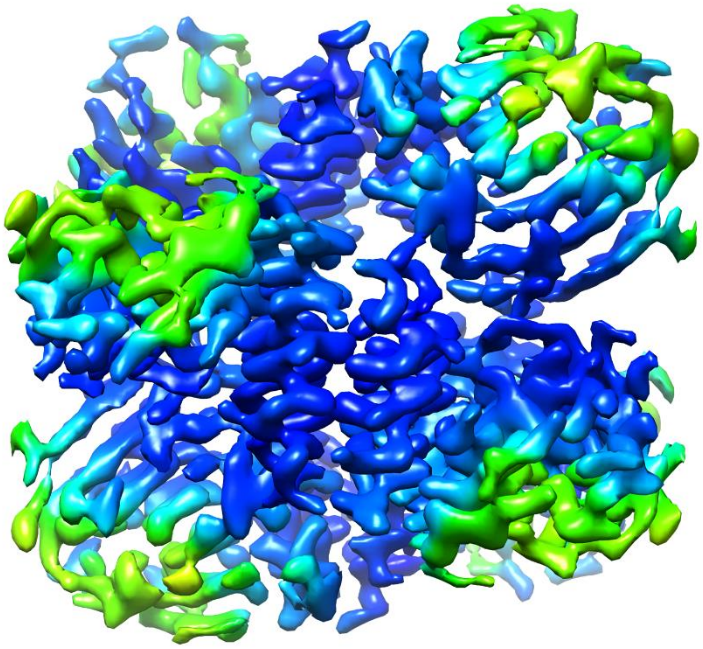

**B**

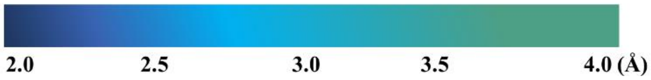

**Figure S3.**

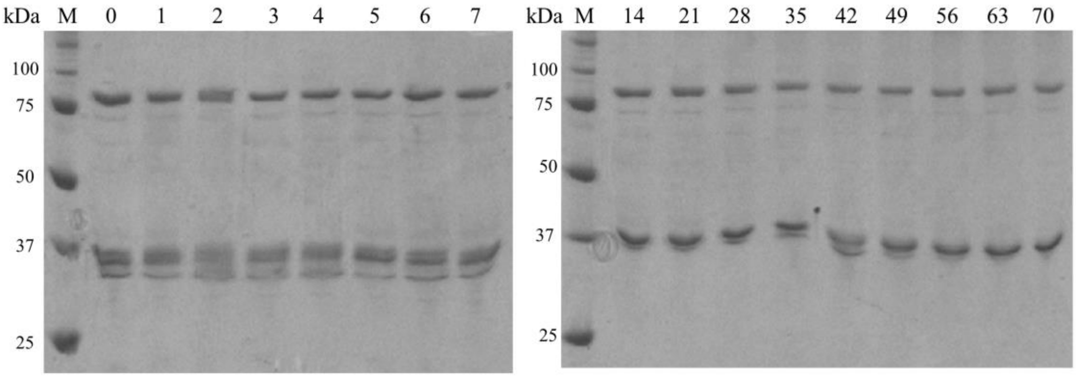


**Figure S4.**

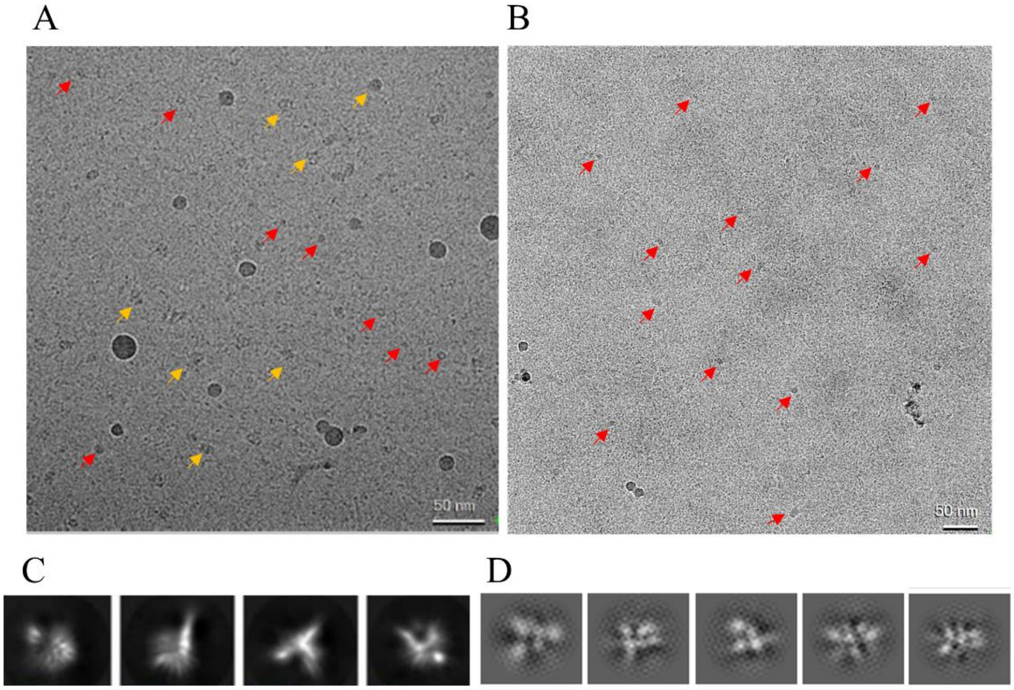


**Table S1.**

| Standard marker | Molecular weight (kDa) |
|---|---|
| Vitamin B12 | 1,350 |
| Myoglobin | 17,000 |
| Ovalbumin | 44,000 |
| Conalbumin | 76,000 |
| Y-globulin | 158,000 |
| Ferritin | 474,000 |
| Thyroglobulin | 660,000 |
| Dextran blue | 2,000,000 |
